# Time of day of infection shapes development of a eukaryotic algal-*Nucleocytoviricota* virocell

**DOI:** 10.1101/2024.05.14.594171

**Authors:** Emily E. Chase, Alexander R. Truchon, Brooke A. Creasey, Steven W. Wilhelm

**Affiliations:** Department of Microbiology, University of Tennessee, Knoxville, TN 37996, United States

**Keywords:** virus-host system, harmful algal bloom, virocell, photosynthesis

## Abstract

*Aureococcus anophagefferens* represents one component of a model host-virus system (with the “giant virus” *Kratosvirus quantuckense*). Studies to define its ribocell (uninfected cells) and virocell (virus-infected cells) forms are needed, as both are abundant during algal blooms. A linkage between light-derived energy, virus particle production and virocell formation has been noted. We explored how the time of day (morning, afternoon, late day) of virus-host contact shaped virocell ontogeny. In parallel, we explored the need for light derived energy in this mixotrophic plankter by inhibiting photosystem II (PSII). Using flow cytometry and photochemical assessments, we examined the physiology of infected cells and controls, and estimated virus particle production by virocells. We observed distinct differences between ribocell and virocell response to treatments, including reductions in virus particle production during reduced light (*i.e.,* duration) and PSII inhibition. Collectively this work demonstrates the importance of light in shaping the fate of infected cells and provides insight into the factors that constrain *in situ* blooms. Most significantly, we show that time of day when a virus and host come into contact influences viral particle production, and therefore bloom dynamics; a factor that needs to be considered in future bloom modeling work.

## INTRODUCTION

The pelagophyte *Aureococcus anophagefferens* was first identified as a harmful algal bloom agent during the formation of brown-tide in Narragansett Bay, Rhode Island in 1985 and described using transmission electron microscopy (Sieburth et al., 1988). As part of that initial study (their Figures 12-16), researchers documented the potential role of infectious virus particles in shaping the fate of cells within the bloom. Subsequent work has commonly focused on the algae in isolation: researchers sequenced the genome of *A. anophagefferens* to provide insight into this phytoplankter’s bloom success through a lens of physiology and environmental interactions (Gobler et al., 2011). More recently, resequencing of the original strain and two additional strains, as well as the assembly of another through publicly available data, offered further insights (Gann et al., 2022). Since then an additional strain has also been made publicly available (Chase et al., 2024). In parallel, a giant virus (*Nucleocytoviricota*) that infects *A. anophagefferens* CCMP1984 and is consistent with the images originally collected by Sieburth and colleagues (1988) has been characterized (Rowe et al., 2008; Brown and Bidle, 2014). Originally known as Aureococcus anophagefferens Virus (AaV), this virus was renamed *Kratosvirus quantuckense* and placed in the family *Schizomimiviridae* by the *International Committee on the Taxonomy of Viruses* (Lefkowitz et al., 2018; Aylward et al., 2023). Recently the genome of *K. quantuckense* has also been re-sequenced and is publicly available (Truchon et al., 2022). The host-virus infection cycle has been explored using transcriptomics following temporal changes from early to late-stage infection in parallel to gene expression of uninfected cells (Moniruzzaman et al., 2018). Cumulatively, the existing characterisation of this system promotes its uses for studying virological concepts in general (Truchon et al., 2023): of particular interest is the virocell concept and the role of cellular energy in virus production.

In the mid-2000s, it was acknowledged that virology disproportionately focused on virus particles without full consideration of infected-cell state; the “virocell” (Forterre, 2011). The importance of the virocell concept was embodied by the ability of a “Mimivirus” (Raoult et al., 2004) to produce viral factories within its host (*i.e.,* functional metabolism and reproduction). Subsequently infected cells were shown to be susceptible to “sickness”, resulting in a reduction of fitness (*e.g.,* reduction of infectious particles) after co-infection of the Sputnik ssDNA virus (La Scola et al., 2008; Desnues et al., 2012). These distinctions (*e.g*., metabolism and reproduction) are absent in most free-floating virions with some exceptions (Moniruzzaman et al., 2018; Gann et al., 2020c). Broadly, when infected cells are transformed to dedicate their metabolic activity to the requirements of the viral genome (Forterre, 2011; Zimmerman et al., 2020) –it has been argued that an infected cell is a cell no more, but instead simply the reproductive phase of the virus (Forterre, 2016). Regardless, virocells require studies separate from their uninfected counterparts and the virion. The viral genome may also introduce auxiliary metabolic genes (AMGs) into these transformed cells, altering the metabolic capabilities of the original cell (Mann et al., 2003; Hurwitz et al., 2013; Vincent and Vardi, 2023). In contrast to the virocell, uninfected cells which rely on encoded ribosomes and focus on the replication of strictly cellular material can be termed “ribocells”. This distinction acknowledges that during early infection both phases are inhabiting the same physical space as the transition from ribocell to virocell occurs (this in between phase has been termed the “ribovirocell” (Forterre, 2011)). This transition has been observed through transcriptional changes during infection (*e.g., Emiliania huxleyi* with Emiliania huxleyi Virus (EhV), and *A. anophagefferens* with *K. quantuckense* (Moniruzzaman et al., 2018; Ku et al., 2020)) and dsRNA-immunofluorescence detection of RNA virus (HcRNAV) infected cells within a population of dinoflagellates (*Heterocapsa circularisquama*) (Coy et al., 2023). Deeper studies have focused on cell physiology during this continuum, primarily looking at cell metabolism.

Cellular metabolism during viral infection takes energy and indeed metabolism is also novel in members of the *Nucleocytoviricota*. As summarised in a recent review (Brahim Belhaouari et al., 2022), *Nucleocytoviricota* collectively encode genes for glycolysis and gluconeogenesis (Moniruzzaman et al., 2020; Ha et al., 2021), fermentation (Schvarcz and Steward, 2018), the Krebs cycle (Moniruzzaman et al., 2020; Ha et al., 2021), lipid metabolism (Nissimov et al., 2019), energy harvesting from inorganic compounds (Schulz et al., 2020), photosynthesis (Moniruzzaman et al., 2017), and reactive oxygen species regulation (Sheyn et al., 2016). Given our limited knowledge of the energetic costs of these metabolic changes during infection by *Nucleocytoviricota,* we sought to characterize light drive energy processes in the virocell of an ecologically important microalgae and its virus, *A. anophagefferens* and *K. quantuckense*.

Here we conducted a series of experiments focused on quantifying photosynthesis in the *K. quantuckense* virocell. It has been acknowledged that, at any given time, an ecologically significant number of cells in marine environments are in an infected state (*i.e.,* virocell) (Suttle and Wilhelm, 1999; Gastrich et al., 2004; Roux et al., 2016; Middelboe and Brussaard, 2017; Carlson et al., 2022). Variations in cellular energetics caused by infection processes would contribute to alterations in ecosystem functions (Middelboe and Brussaard, 2017). Previous work has already shown the importance of light availability during infection of *A. anophagefferens*, where lower light levels led to a reduced burst sizes (Gann et al., 2020); providing insight into bloom dynamics given the occurrence of self-shading in *A. anophagefferens* blooms as well as mixing through the water column. We furthered these studies by focusing on the time of infection (*i.e.,* when a ribocells transitions to a virocell) in relation to virus particle production, and the reliance of the development of viral infection of *A. anophagefferens* on continued photosynthesis by the virocell. This is completed by (1) staggering virocell initiation during a standard light cycle and profiling, and (2) gauging the reaction of ribocells and virocells to a photosynthesis blocker (3-(3,4-dichlorophenyl)-1,1-dimethylurea; DCMU).

## MATERIALS AND METHODS

### Cell culturing and virus propagation

*Aureococcus anophagefferens* (CCMP 1984, Bigelow NCMA) cells were grown on a 12/12 light:dark (∼100 μmol photons m^-2^ s^-1^) cycle at 19° C in ASP12A growth medium (Gann, 2016) as xenic cultures. Additional cultures for virus propagation were grown (∼1 L) as described and inoculated with viral lysate. Upon full lysis (∼one week) infected cultures were filtered through 47-mm diameter, 1.0-μm nominal pore-size Whatman^™^ Nucleopore Track Etch Membrane (Whatman, Inc.) followed by 47-mm diameter, 0.45-μm pore-size Millipore Isopore PC Membrane (MilliporeSigma) to remove cells. The resulting lysate was concentrated on a Tangential Flow Filtration System (Fisher Scientific) with a Durapore 30 kDa Pellicon XL Filter (MiliporeSigma) to a final volume of ∼50 mL for experimental use.

### Procedures for enumerating viral particles and host cells

Viral particle abundance within concentrated lysate (Chase et al., 2022c) and host cell density (Chase et al., 2022d) were both determined by flow cytometry. Briefly, a Cytoflex Flow Cytometer system (Beckman Coulter) equipped with a violet laser (Zhao et al., 2023) was used to enumerate viral particles stained with SYBR Gold Nucleic Acid Stain (Invitrogen). *A. anophagefferens* cells were enumerated based on autofluorescence. The flow cytometer provided additional data for the host cells including relative cell size (violet side scatter; Violet SSC) and relative per-cell autofluorescence (peridinin-chlorophyll protein complex; PerCP). Cell and virus cytometry gating were confirmed with negative controls (media; see **Supplemental Figure 1**). Cell (*i.e.,* ribocells and virocells) photosynthesis (*i.e.,* photochemistry) was defined by pulse amplitude modulation using a Walz Phyto-PAM analyzer Compact Version (Heinz Walz GmbH) as previously described (Chase et al., 2022b) to record F_v_/F_m_, which provides insight into the overall function of photosystem II (*i.e.,* “stress”) (Campbell et al., 1998). *Virocell formation at different times during the solar day*

Experiments on the timing of infection within the solar day were completed in batch cultures with starting cell densities of ∼1.0×10^6^ mL^-1^. Treatments included no-virus negative controls (*i.e.,* ribocells) and treatments inoculated with viruses at three different times in the solar day: infections occurred with 12 h (EARLY-V), 6 h (MID-V), and 20 min (LATE-V) time remaining exposure to light. All experiments were conducted in true biological triplicate.

Cultures were infected at a multiplicity of infection (MOI) of approximately 75 particles per cell. We note that MOI here reflects the total abundance of viral particles administered and not number of infectious units. Samples for flow cytometry (*i.e.,* cell population, relative size, and PerCP) were collected for 74 h across 14 timepoints. For each of infection treatments virus abundance was also measured. Virus sampling occurred at 0, 6, 12, 24, 24.5, 30, 48, and 72 hours post infection (hpi) relative to the initial time point of the EARLY-V infection. See **Supplemental Figure 2** for an experiment schematic. For virus enumeration at 24, 48, and 72 hpi (start of light cycle), cultures were permitted approximately 20 min in light before samples were taken for virus staining, this maximises the number of produced viruses being captured in our counts by giving time for lysis to occur (this practice was carried out for all experiments).

### Inhibition of photosystem II by DCMU

DCMU (3-(3,4-dichlorophenyl)-1,1-dimethylurea; *aka* diuron) (40 μM final concentration) was applied to cells to inhibit PSII at the start of their light cycle. For this component batch cultures at cell concentrations of ∼3.0×10^6^ mL^-1^ were prepared: treatments included a no DCMU/no virus control (*i.e.,* ribocells), a no virus control (with DCMU), a virus treatment (no DCMU, *i.e.,* virocells), and a DCMU plus viral inoculated treatment. All experiments were completed in triplicate. Virus inoculated treatments were infected at a MOI of approximately 42 particles per cell. DCMU and virus treatments were permitted to acclimate to DCMU ∼10 min in light before the administering of the viral inoculant. This application time coincides with the time required for the PSII to be inhibited by DCMU in the algae *Tetraselmis spp.* (Kristoffersen et al., 2016). DCMU concentration was determined by a preliminary experimentation observing both cells counts (**Supplemental Figure 3**) and F_v_/F_m_(**Supplemental Figure 4**). Flow cytometry measurements were collected at ten times across 49 hours post infection (hpi) and post-DCMU application. Viral particles were enumerated at 0, 2, 24, 26, 49 hpi. Samples for photochemical measurements occurred at 0, 2, 11.5, 24, 26, 28, and 48 hpi and were dark-adapted for 30 min before analysis at a low light level (gain = 13, photosynthetically active radiation (PAR) = 1, λ = 440 nm). The experiment took place over two light cycles.

### Statistical Analyses

Data management, analyses, and visualisation were carried out using R Statistical Software (v22.12.0; (R Core Team, 2021)) by packages *tidyr* (Wickham et al., 2023), *tidyverse* (Wickham et al., 2019), *dplyr* (Wickham et al., 2022), *stringr* (Wickham, 2023), *ggplot2* (Wickham, 2011). Statistical tests (Welch’s t-tests) were run using base R for comparisons within experiments. For all tests we assumed used a threshold of p < 0.05 for significance, but we provide these numbers so the reader can decide. Growth rates were calculated as the change in net cell population counts (as measured by cytometry) over two day/night cycles (48 h).

## RESULTS

### Virus particle production increases with a longer light exposure

After two day:night cycles (48 hpi), the abundances of virus particles produced were lower in virocells exposed to shorter light periods (**Figure 1A**, **Supplemental Figure 5**). Virus lysate addition late in the solar day (*i.e.,* LATE-V) produced fewer viral particles than those infected at the beginning (12 h exposure; EARLY-V) and middle of the light cycle (6 h exposure; MID-V) (p = 0.072 (early *vs* mid) and p = 0.008 (early *vs* late), **Supplemental Table 1**). After three light cycles (72 h), there was a difference between the abundance of viral particles produced between the mid to late samples (p = 0.048), but no longer between late and early (p = 0.108), seemingly the effect of virocells being formed later in a light cycle is reduced over additional light cycles using a relatively high MOI (75). Cell lysis was also occurring after 24, 48, and 72 hpi (**Supplemental Figure 6**, **Supplemental Table 2**). Relative cell size (by violet side scatter) incrementally increased in all treatments except for the control where cell division throughout the night “reset” the population’s mean cell size, conversely virocells continued to enlarge (although we recognise that populations exposed to lysate would still include some ribocells—which may harbour infection resistance, we assume it is primarily made up of virocells given these changes in cytometry measurements) (**Figure 1B**, **Supplemental Figure 7**, **Supplemental Table 2**). A similar pattern as relative cell size occurs with PerCP (**Supplemental Figure** 8). At 24, 48, and 72 hpi (*i.e.,* one, two, and three complete light cycles) the effects of even the late viral lysate application were apparent in relative size of *A. anophagefferens*, where cells infected ∼20 min before the dark cycle were smaller than both the “early” and “mid” infections (early *vs.* mid for 24, 48, and 72 hpi respectively: p-values: 0.017 and 0.024, < 0.001 and 0.022, < 0.001 and 0.016).

**Figure 1.**
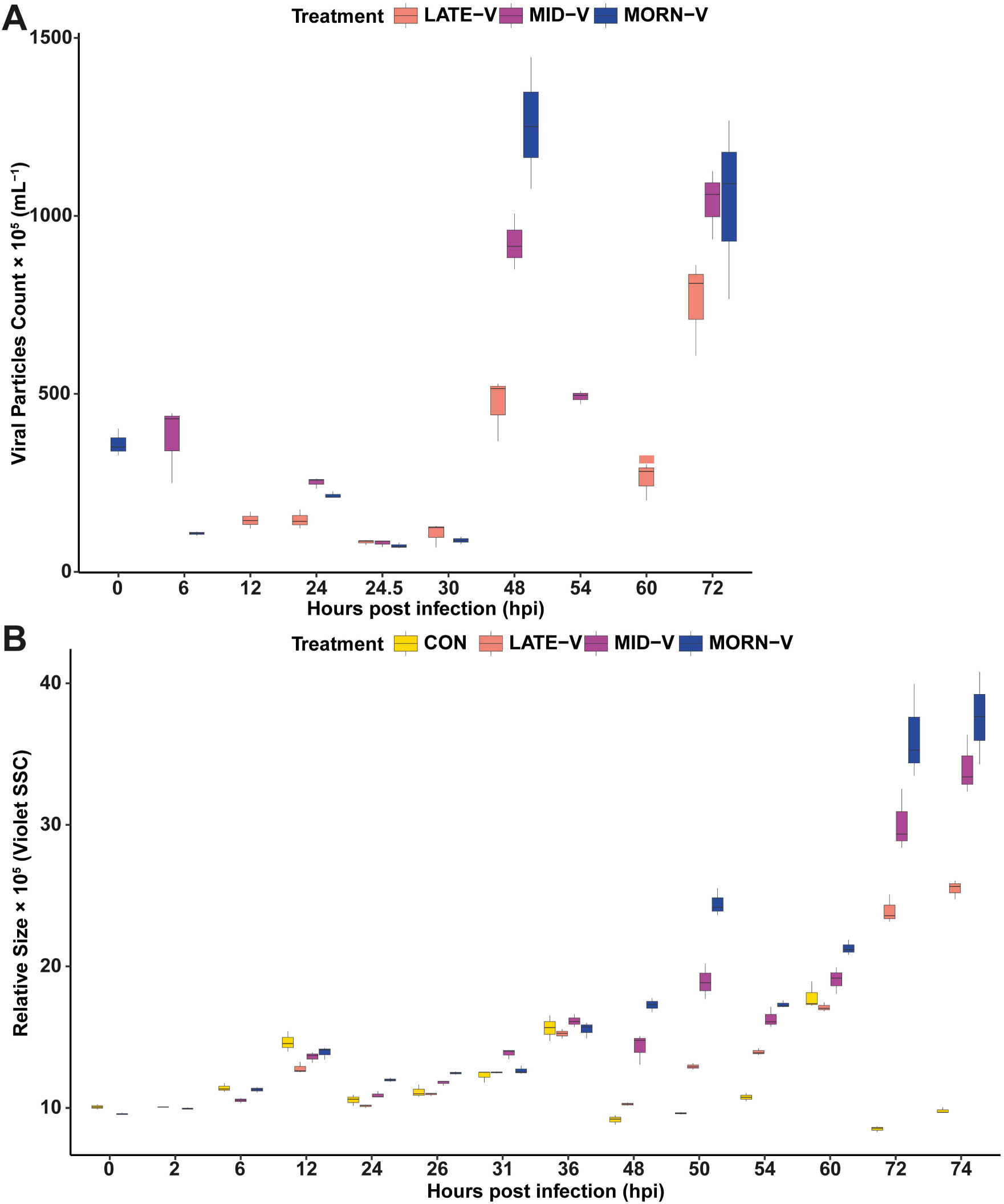
Virocell formation (*i.e.,* virus lysate application) at different time points within a 12/12 light/dark cycle in *Aureococcus anophagefferens* with the virus *Kratosvirus quantuckense*. **(A)** Mean viral particle counts (within culture media) at different time points (all relative to the early infection start time). **(B)** Mean relative size of *A. anophagefferens* population cells throughout three light cycles (plus two hours after the third light cycle) as measured via cytometry. Abbreviations: LATE-V; lysate applied ∼20 min before the end of the 12 h light cycle, MID-V; lysate applied midway through the light cycle (6 h), EARLY-V; lysate applied at the start of the light cycle (12 h exposure), CON; ribocell control (no viral lysate); SSC: side scatter. **Alt text:** A boxplot shows that the number of viral particles produced after forty-eight experimental hours and seventy-two experimental hours is more in cell infected early in the day than late in the day. A second boxplot shows that the average cell size of a population at seventy-two hours is larger in cells infected early in the day than later in the day.

### Interruption of PSII electron transport inhibits population growth and virus production

For *A. anophagefferens* cultures acclimated to a 12/12h light/dark cycle the application of DCMU halted photosynthesis within 40 min (when the first photochemistry analyses occurred) (**Figure 2A**, **Supplemental Figure 9**) as indicated by a statistically significant difference between F_v_/F_m_ of the ribocells without DCMU (*i.e.,* control) and DCMU applied ribocells (p= < 0.000, **Supplemental Table 3**), a trend that continued at each sampling point. These results are unsurprising given the use of DCMU to block photosynthesis in other eukaryotic algae (*e.g.,* (Gonen-Zurgil et al., 1996; Camuel et al., 2017)) and for an alternate photosynthetic efficiency calculation in *A. anophagefferens* (Gobler et al., 2007). Additionally there was also a “DCMU enhanced fluorescence” response in photochemistry inferred relative chlorophyll content as previously shown in *A. anophagefferens* (Keller and Rice, 1990), but only for two hours post infection as it was no longer present at 11.5 h into the DCMU application (**Figure 2B**, **Supplemental Figure 10**, **Supplemental Table 4**; at 0 hpi, p = 0.000; at 2 hpi, p = 0.000, at 11.5 hpi p = 0.860). We note we did not see this pattern in our cytometry PerCP (peridinin-chlorophyll protein complex) data (**Supplemental Figure 10**).

**Figure 2.**
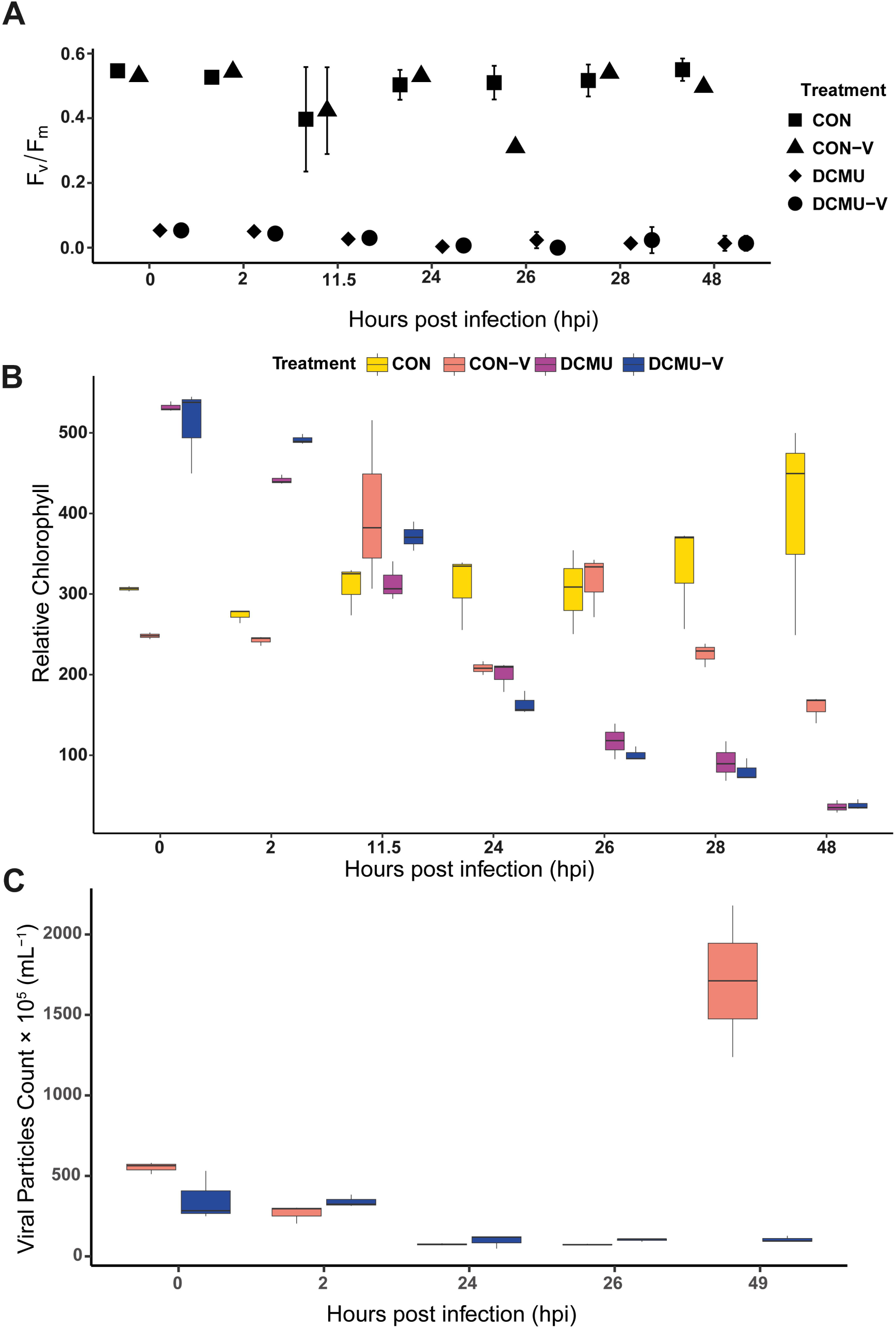
*Aureococcus anophagefferens* response to application of 40 μM DCMU (3-(3,4-dichlorophenyl)-1,1-dimethylurea; diuron) and infection by *Kratosvirus quantuckense*. **(A)** Mean maximum quantum yield of PSII at a low light level (PAR = 1) over two light cycles. **(B)** Relative chlorophyll (photochemistry) over two light cycles**. (C)** In media viral particle count throughout the infection. Abbreviations: CON; control (ribocells), CON-V; virocells without DCMU applied, DCMU; ribocells with DCMU applied, DCMU-V; virocells with DCMU applied. **Alt text:** A boxplot shows that the photosynthetic efficiency of infected and uninfected cells exposed to DCMU is virtually zero while the photosynthetic efficiency of cells not exposed to DCMU is at about 60%. A boxplot shows that the relative chlorophyll content after forty-eight experimental hours is much higher in uninfected cells with DCMU exposure than both infected and uninfected cells with DCMU exposure, and then infected cells without DCMU exposure. A boxplot showing that after forty-nine experimental hours the number of viral particles produced is virtually zero in infected cells exposed to DCMU and high in infected cells without DCMU exposure.

Cytometry based cell counts between control ribocells and DCMU exposed ribocells (no virus) were different at 24, and 29 hpi (p = 0.016, and 0.006, respectively), and a similar pattern was shown at 26 and 48 hpi (p = 0.052 and 0.052; **Supplemental Figure 11**, **Supplemental Table 5**). As expected, at 48 hpi there was a significant difference between the control and virus-infected populations (both with and without DCMU; p = 0.038 and 0.047). There were more cells in DCMU treated control cultures than the virus infected (*i.e.,* virocells) non-DCMU cultures (p = 0.019). There was no difference between cultures treated only with DCMU and cultures treated with DCMU and viral lysate (p = 0.169, **Supplementary Table 5**).

There was a difference (albeit non-significant) in growth rate at 48 h of the control and the DCMU treated ribocells (**Supplemental Figure 12**; **Supplemental Table 6**, p = 0.056). Given the growth rate for the DCMU-treated cells was negative, it is possible the DCMU population would collapse if given additional light cycles as DCMU causes a non-reversible block of electron transfer (Kirilovsky et al., 1994). Previous work has shown that DCMU protects the D1 protein from degradation and removal (Chung and Jung, 1995) and only under “high light” conditions will DCMU fail to protect against photodamage—this “high light” level required in our system is yet to be known and outside the scope of this study. Nonetheless, DCMU was effective at blocking PSII. Growth rate was also not different (p = 0.184) between cultures with virus lysate applied independent of the presence of DCMU. Cultures acclimated to DCMU and with viral lysate had fewer free viral particles recorded after two light cycles (48 hpi) than infected cultures without DCMU (**Supplemental Table 7**, p = 0.027) (**Figure 2C, Supplemental Figure 13**). Finally, relative size (by violet side scatter) showed the same pattern as discussed for the “light exposure” experiment (**Supplemental Figure 14**).

## DISCUSSION

We applied treatments to the *A. anophagefferens* and *K. quantuckense* host-virus system to assess responses to light derived energy during virus infection and production. Our results present a unique perspective in that the duration of light exposure to the host-virus pair after infection has implications for the abundance of viral particles being released at the time of lysis. This suggests that the time of day that a virus contacts and infects a host may be critical to the outcome of that encounter. Additionally, we provide insight into the complete disruption of photosynthetic energy production by inhibition of the PSII D1 protein using DCMU, showcasing that infection in absence of photosynthesis (akin to no daylight) also produces few to no viral particles.

### The infection cycle of A. anophagefferens by K. quantuckense is constrained by the length of light exposure and occurance of a “dark period”, not by time since virus contact/infection

A major observation of these studies is that the “standard” infection cycle of *A. anophagefferens* with *K. quantuckense* (reported as ∼24 h during previous experiments, (Brown and Bidle, 2014; Moniruzzaman et al., 2018; Gann et al., 2020b)) is reflective of the light/dark cycles used during experiments in laboratories. Our results demonstrate that the time from infection to lysis is influenced by cumulative light and dark periods, rather than time since infection, although some theoretical maximum likely occurs based on cell space for viral particles within the host cell, *etc*. For cultures exposed to the same irradiance time and on the same light cycle (*i.e.,* acclimated to the same growing conditions), we demonstrated that the length of the light period after virus contact had consequences for the abundance of viral particles produced—even after several light/dark cycles. This observation demonstrates important lasting consequences (at least over three “days”) of timing of the virus-host contact (relative to the dark cycle) on infection dynamics. Previous work has found that infection cycle length can change among cells exposed to differing light intensity (see (Gann et al., 2020a; Gobler et al., 2007). Consequently, there is an interplay between light availability (intensity) and exposure period with virus production in this system. In natural systems where *A. anophagefferens* blooms (like Quantuck Bay, NY) the difference in solar day length can be as much as 2 h per day (June (15h) relative to late August (13h)) during the summer bloom season. While we did not test these specific times in the current study, based on our observations the reduction in day length of ∼13% during the bloom season may potentially have effects on virus-host interactions.

The combined roles of light and time of viral contact are likely an important factor in bloom termination/collapse among *in situ A. anophagefferens* and any *K. quantuckense* or *K. quantuckense*-related viruses (Moniruzzaman et al., 2017), especially as irradiance is not homogenous among the population—which is specifically relevant to the turbid bays where *A. anophagefferens* blooms. Daylength (alongside nutrients, temperature, *etc*.) may also contribute to the duration of *A. anophagefferens* blooms, which can “peak” for a span of 20 days (Wazniak and Glibert, 2004) or produce multiple bloom peaks in a season (Moniruzzaman et al., 2016). At this point we note this is a study of just one virus-host system alone, and that *in situ* other *K. quantuckense*-related viruses may exhibit different energetics (*e.g.,* physiology, burst sizes, viral particle production in relation to light, *etc.*). In terms of evolution, we relate this to the so-called “quasigenus” effect, where giant viruses diversify into a closely related group capable of different infection strategies to match different strains of a bloom species (*e.g., A. anophagefferens*) and environmental (including light) conditions (Highfield et al., 2014, 2017; Chase et al., 2022a). Seemingly, the mixotrophic capabilities of *A. anophagefferens* should help the host persist in a bloom state during light-limitation, with the consequence of reducing viral production. This does lead to an interesting survival strategy for the host: switching to heterotrophic growth may provide an extended period for host-defense mechanisms to purge an infectious virus.

### Viral particle production within PSII blocked cells: questioning mixotrophy

*A. anophagefferens* cultures acclimated to DCMU produced significantly fewer virus particles compared to cultures not treated with DCMU, implying light is necessary for viral particle production. This is also evident in some cyanobacterium-bacteriophage systems (Ni and Zeng, 2016; Liu et al., 2019). We note that cell acclimation to DCMU was carried out during the light cycle (*i.e.,* treatment was applied near the start of the light cycle but not in the complete absence of light). Despite our cells’ response to DCMU (*e.g.,* reduced growth rates), it is not definite that DCMU application would necessarily increase the progression of photodamage in the presence of light and consequently cell death (Kirilovsky et al., 1994; Allakhverdiev et al., 2005). In fact, algae capable of mixotrophy are potentially resistant to DCMU as an algaecide (and “herbicides” in general (Xie et al., 2023)). *A. anophagefferens* is capable of mixotrophy (Cosper et al., 1989; Dzurica et al., 1989; Gobler and Sañudo-Wilhelmy, 2001), so this disconnect requires further exploration given that cells grown under mixotrophic conditions could respond differently. In theory, blocking PSII of *A. anophagefferens* and forcing them into a heterotrophic lifestyle should give insights into viral production driven by heterotrophic energy production: as noted above that there is little virus production this may be part of a larger resistance mechanism in these cells. In the present case, where PSII was inhibited at the start of the light cycle, cells produced fewer viral particles than cells which did not have PSII interrupted. These results provide further evidence of the importance of light to virus infection.

### Initial light period and exposure shapes infection dynamics

Incorporation of light history and irradiance along with the timing of virus-host contact within the light cycle should be considered in studies of *A. anophagefferens* and *K. quantuckense*. In the present study we attempted to mimic a “time of day” infection model – contrasting virus-host infections that would happen early in the solar day *vs* later. This is perhaps most relevant (and complicating) for *in situ* studies where cells in a population are metabolically heterogenous (*e.g.,* employing photosynthesis *vs.* heterotrophic strategies to acquire energy) due to varying light availabilities and histories. Based on our conclusions, infection dynamics should differ based on when a virus encounters a cell during the solar day (relative to nightfall), the amount of light attenuation in the water column, bloom heterogeneity and mixing, bloom density (self-shading), and how photosynthesis in general is proceeding in individual cells throughout the population (as demonstrated by our DCMU experiment) (see Featured Figure, **Figure 3**), and additionally the sinking rate of cells which we know differs between infected and uninfected cells (Truchon et al., 2024).Going forward, these factors need to be considered for *in situ* bloom models.

**Figure 3.**
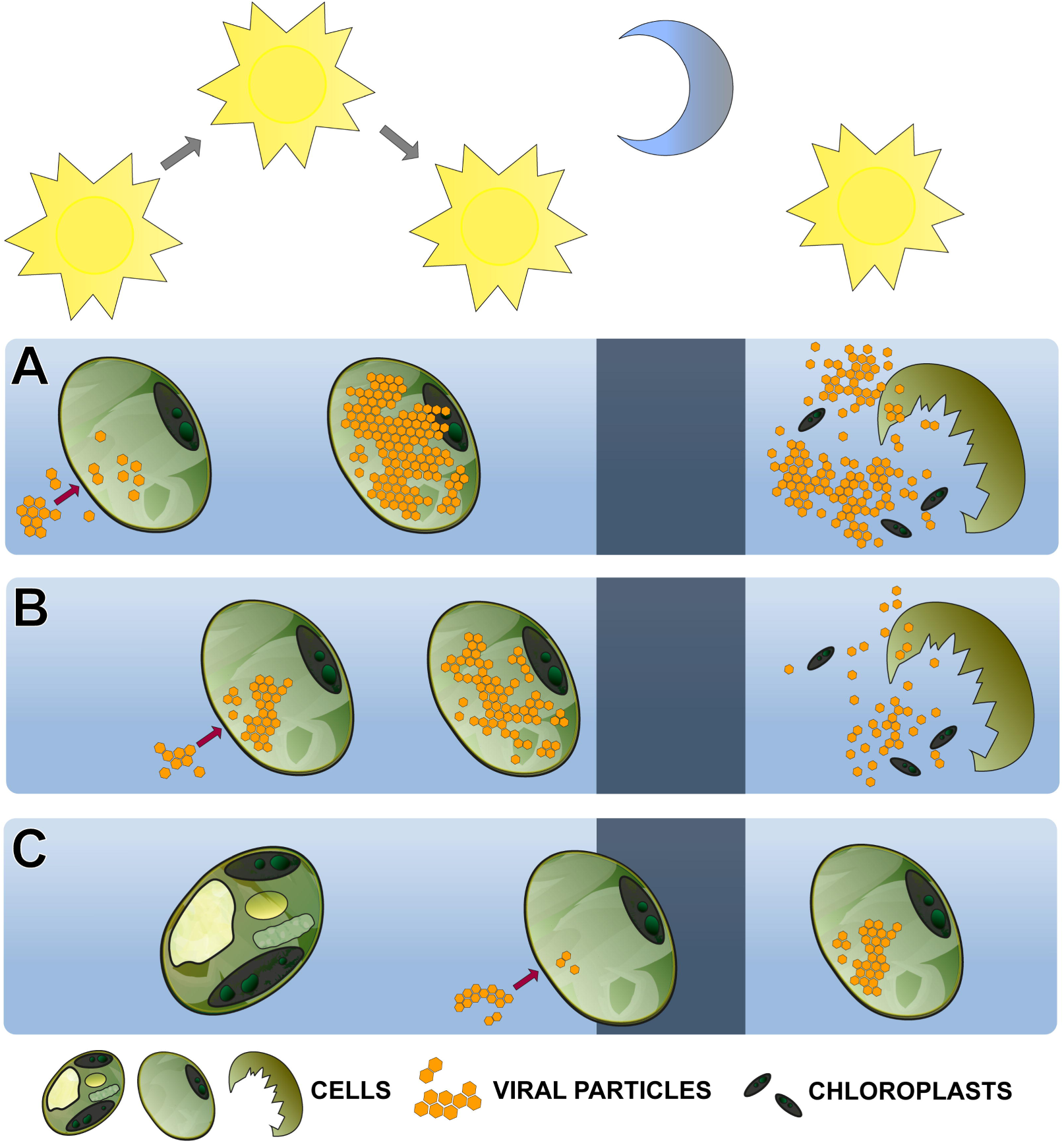
Conclusive schematic of the collective results of these studies. (**A**) Cell populations infected early in the day (longest light exposure) produce many viral particles at the time of lysis early the next day, while (**B**) populations infected middle of the day produce fewer viral particles, and (**C**) populations infected near the end of the day produce the least number of viral particles. Chloroplasts remain intact during the infection cycle in situations **A-B**. **Alt text:** A cartoon depicting *Aureococcus anophagefferens* and the production of its virus, *Kratosvirus quantuckense*. The image shows infection at different times of day and how infection later in the day produces fewer viral particles than early in the day.

In the context of *A. anophagefferens*, which is known to bloom in coastal bays (Mulholland et al., 2004; Sieracki et al., 2004; Simjouw et al., 2004; Probyn et al., 2010; Yao et al., 2019), the effects of climate change (*e.g.,* temperature, nutrient composition, mixing and water column structure) can shape light availability. An indirect consequence of these changes may be altered bloom duration due to light constraints on virus-mediated bloom termination. Decreasing virus production and cell lysis could lead to enhanced light attenuation and a positive feedback loop that could increase bloom duration and affect the local ecosystem. Consequently, light histories become important metrics (similar to nutrients, temperature, *etc.*) that need to be considered and if possible, hindcast for environment-based experiments.

## CONCLUSION

We explored the effects of “time of day” of virus-host contact, and inhibition of photosynthesis on virus production in an ecologically important HAB-giant virus system. Our results also showcase how within the same algal species ribocells and virocells (both important and significant components of a population) differ, and that bloom heterogeneity needs to be considered. A better understanding of the role of viruses in the ecology of blooms of this brown tide causative agent will require an understanding of light history (intensity and time), and heterogeneity (ribocell vs virocell) to provide both a better mechanistic understanding of the potential role of viruses and cell state in bloom demise.

## AUTHOR CONTRIBUTIONS

**EEC:** conceptualization, formal analysis, investigation, methodology, visualization, writing – original draft preparation, project administration. **ART:** writing – review & editing. **BAC:** methodology**. SWW:** Conceptualisation, funding acquisition, project administration, supervision, validation, writing – review & editing

## ACKNOWLEDGEMENTS

The authors wish to thank Brittany N. Zepernick, Katelyn A. Houghton, and Gwen F. Stark for their feedback and discussion on these methods and conclusions.

---

**Supplemental File 1.** Composition of additional figures showing methodology, experimental schematic, and cytometry and photochemistry results. Also includes tables presenting statistical analyses.

## REFERENCES

Allakhverdiev, S. I., Nishiyama, Y., Takahashi, S., Miyairi, S., Suzuki, I., and Murata, N. (2005). Systematic analysis of the relation of electron transport and ATP Synthesis to the photodamage and repair of photosystem II in *Synechocystis*. Plant Physiology 137, 263–273. doi: 10.1104/pp.104.054478

Aylward, F. O., Abrahão, J. S., Brussaard, C. P. D., Fischer, M. G., Moniruzzaman, M., Ogata, H., et al. (2023). Taxonomic update for giant viruses in the order Imitervirales (phylum *Nucleocytoviricota*). Archives of Virology 168, 283. doi: 10.1007/s00705-023-05906-3

Brahim Belhaouari, D., Pires De Souza, G. A., Lamb, D. C., Kelly, S. L., Goldstone, J. V., Stegeman, J. J., et al. (2022). Metabolic arsenal of giant viruses: Host hijack or self-use? eLife 11, e78674. doi: 10.7554/eLife.78674

Brown, C. M., and Bidle, K. D. (2014). Attenuation of virus production at high multiplicities of infection in *Aureococcus anophagefferens*. Virology 466–467, 71–81. doi: 10.1016/j.virol.2014.07.023

Campbell, D., Hurry, V., Clarke, A. K., Gustafsson, P., and Öquist, G. (1998). Chlorophyll fluorescence analysis of cyanobacterial photosynthesis and acclimation. Microbiology and Molecular Biology Reviews 62, 667–683. doi: 10.1128/mmbr.62.3.667-683.1998

Camuel, A., Guieysse, B., Alcántara, C., and Béchet, Q. (2017). Fast algal eco-toxicity assessment: Influence of light intensity and exposure time on *Chlorella vulgaris* inhibition by atrazine and DCMU. Ecotoxicology and Environmental Safety 140, 141–147. doi: 10.1016/j.ecoenv.2017.02.013

Carlson, M. C. G., Ribalet, F., Maidanik, I., Durham, B. P., Hulata, Y., Ferrón, S., et al. (2022). Viruses affect picocyanobacterial abundance and biogeography in the North Pacific Ocean. Nature Microbiology 7, 570–580. doi: 10.1038/s41564-022-01088-x

Chase, E. E., Pitot, T. M., Monteil-Bouchard, S., Desnues, C., and Blanc, G. (2022a). “Quasigenus” among Phycodnaviridae: A diversity of chlorophyte-infecting viruses in response to a dense algal culture in a high-rate algal pond. 2022.12.26.521929. doi: 10.1101/2022.12.26.521929

Chase, E. E., Stark, G., Truchon, A., and Wilhelm, S. W. (2022b). Application of PHYTO-PAM-II (Compact Version) on Aureococcus anophagefferens cultures for photosynthetic efficiency and quantum yield of PSII. Available at: https://www.protocols.io/view/application-of-phyto-pam-ii-compact-version-on-aur-cgt4twqw (Accessed April 25, 2023).

Chase, E. E., Truchon, A., Coy, S. R., and Wilhelm, S. W. (2022c). Aureococcus anophagefferens Virus (AaV)/Kratosvirus quantuckense viral particle count by SYBR Green staining and flow cytometry (CytoFLEX S Flow Cytometer Beckman Coulter) Available at: https://www.protocols.io/view/aureococcus-anophagefferens-virus-aav-floreovirus-ci99uh96 (Accessed April 25, 2023).

Chase, E. E., Truchon, A. R., Schepens, W. W., and Wilhelm, S. W. (2024). *Aureococcus anophagefferens* strain CCMP1851: Draft genome of a second *Kratosvirus quantuckense* susceptible host strain for this host-giant virus model system. Microbiol Resour Announc. doi:10.1128/mra.00292-24

Chase, E. E., Truchon, A., and Wilhelm, S. W. (2022d). Aureococcus anophagefferens population count, and relative size (Violet SSC) by flow cytometry (CytoFLEX S Flow Cytometer Beckman Coulter). Available at: https://www.protocols.io/view/aureococcus-anophagefferens-population-count-and-r-cgzftx3n (Accessed April 25, 2023).

Chung, S. K., and Jung, J. (1995). Inactivation of the acceptor side and degradation of the D1 protein of photosystem II by singlet oxygen photogenerated from the outside. Photochemistry and Photobiology 61, 383–389. doi: 10.1111/j.1751-1097.1995.tb08627.x

Cosper, E. M., Dennison, W., Milligan, A., Carpenter, E. J., Lee, C., Holzapfel, J., et al. (1989). An examination of the environmental factors important to initiating and sustaining “brown tide” blooms., in Novel Phytoplankton Blooms, eds. E. M. Cosper, V. M. Bricelj, and E. J. Carpenter (Berlin, Heidelberg: Springer), 317–340. doi: 10.1007/978-3-642-75280-3_18

Coy, S. R., Utama, B., Spurlin, J. W., Kim, J. G., Deshmukh, H., Lwigale, P., et al. (2023). Visualization of RNA virus infection in a marine protist with a universal biomarker. Sci Rep 13, 5813. doi: 10.1038/s41598-023-31507-w

Desnues, C., Boyer, M., and Raoult, D. (2012). “Chapter 3 - Sputnik, a virophage infecting the viral domain of life,” in Advances in Virus Research, eds. M. Łobocka and W. T. Szybalski (Academic Press), 63–89. doi: 10.1016/B978-0-12-394621-8.00013-3

Dzurica, S., Lee, C., Cosper, E. M., and Carpenter, E. J. (1989). Role of environmental variables, specifically organic compounds and micronutrients, in the growth of the Chrysophyte *Aureococcus anophagefferens*. In Novel Phytoplankton Blooms E. M. Cosper, V. M. Bricelj, and E. J. Carpenter (Berlin, Heidelberg: Springer), 229–252. doi: 10.1007/978-3-642-75280-3_13

Forterre, P. (2011). Manipulation of cellular syntheses and the nature of viruses: The virocell concept. Comptes Rendus Chimie 14, 392–399. doi: 10.1016/j.crci.2010.06.007

Forterre, P. (2016). To be or not to be alive: How recent discoveries challenge the traditional definitions of viruses and life. Studies in History and Philosophy of Science Part C: Studies in History and Philosophy of Biological and Biomedical Sciences 59, 100–108. doi: 10.1016/j.shpsc.2016.02.013

Gann, E. R. (2016). ASP12A Recipe for culturing *Aureococcus anophagefferens*. protocols.io. doi: dx.doi.org/10.17504/protocols.io.f3ybqpw

Gann, E. R., Gainer, P. J., Reynolds, T. B., and Wilhelm, S. W. (2020a). Influence of light on the infection of *Aureococcus anophagefferens* CCMP 1984 by a “giant virus.” PLOS ONE 15, e0226758. doi: 10.1371/journal.pone.0226758

Gann, E. R., Hughes, B. J., Reynolds, T. B., and Wilhelm, S. W. (2020b). Internal Nitrogen Pools Shape the Infection of *Aureococcus anophagefferens* CCMP 1984 by a Giant Virus. Frontiers in Microbiology 11. Available at: https://www.frontiersin.org/articles/10.3389/fmicb.2020.00492 (Accessed July 11, 2023).

Gann, E. R., Truchon, A. R., Papoulis, S. E., Dyhrman, S. T., Gobler, C. J., and Wilhelm, S. W. (2022). *Aureococcus anophagefferens* (Pelagophyceae) genomes improve evaluation of nutrient acquisition strategies involved in brown tide dynamics. Journal of Phycology 58, 146–160. doi: 10.1111/jpy.13221

Gann, E. R., Xian, Y., Abraham, P. E., Hettich, R. L., Reynolds, T. B., Xiao, C., et al. (2020c). Structural and proteomic studies of the Aureococcus anophagefferens Virus demonstrate a global distribution of virus-encoded carbohydrate processing. Frontiers in Microbiology 11. doi: 10.3389/fmicb.2020.02047

Gastrich, M. D., Leigh-Bell, J. A., Gobler, C. J., Roger Anderson, O., Wilhelm, S. W., and Bryan, M. (2004). Viruses as potential regulators of regional brown tide blooms caused by the alga, *Aureococcus anophagefferens*. Estuaries 27, 112–119. doi: 10.1007/BF02803565

Gobler, C. J., Anderson, O.R., Gastrich, M., and Wilhelm, S. W. (2007). Ecological aspects of viral infection and lysis in the harmful brown tide alga *Aureococcus anophagefferens*. Aquatic Microbial Ecology 47, 25–36. doi: 10.3354/ame047025

Gobler, C. J., Berry, D. L., Dyhrman, S. T., Wilhelm, S. W., Salamov, A., Lobanov, A. V., et al. (2011). Niche of harmful alga *Aureococcus anophagefferens* revealed through ecogenomics. Proceedings of the National Academy of Sciences 108, 4352–4357. doi: 10.1073/pnas.1016106108

Gobler, C., and Sañudo-Wilhelmy, S. (2001). Effects of organic carbon, organic nitrogen, inorganic nutrients, and iron additions on the growth of phytoplankton and bacteria during a brown tide bloom. Marine Ecology-progress Series - MAR ECOL-PROGR SER 209, 19–34. doi: 10.3354/meps209019

Gonen-Zurgil, Y., Carmeli-Schwartz, Y., and Sukenik, A. (1996). Selective effect of the herbicide DCMU on unicellular algae — a potential tool to maintain monoalgal mass culture of *Nannochloropsis*. J Appl Phycol 8, 415–419. doi: 10.1007/BF02178586

Ha, A. D., Moniruzzaman, M., and Aylward, F. O. (2021). High transcriptional activity and diverse functional repertoires of hundreds of giant viruses in a coastal marine system. mSystems 6, e00293–21. doi: 10.1128/mSystems.00293-21

Highfield, A., Evans, C., Walne, A., Miller, P. I., and Schroeder, D. C. (2014). How many *Coccolithovirus* genotypes does it take to terminate an *Emiliania huxleyi* bloom? Virology 466–467, 138–145. doi: 10.1016/j.virol.2014.07.017

Highfield, A., Joint, I., Gilbert, J. A., Crawfurd, K. J., and Schroeder, D. C. (2017). Change in *Emiliania huxleyi* virus assemblage diversity but not in host genetic composition during an ocean acidification mesocosm experiment. Viruses 9, 41. doi: 10.3390/v9030041

Hurwitz, B. L., Hallam, S. J., and Sullivan, M. B. (2013). Metabolic reprogramming by viruses in the sunlit and dark ocean. Genome Biology 14, R123. doi: 10.1186/gb-2013-14-11-r123

Keller, A. A., and Rice, R. L. (1990). Variation in DCMU-enhanced fluorescence relative to chlorophyll a: Correlation with the brown tide bloom. Journal of Phycology 26, 202–205. doi: 10.1111/j.0022-3646.1990.00202.x

Kirilovsky, D., Rutherford, A. W., and Etienne, A. L. (1994). Influence of DCMU and ferricyanide on photodamage in photosystem II. Biochemistry 33, 3087–3095. doi: 10.1021/bi00176a043

Kristoffersen, A. S., Hamre, B., Frette, Ø., and Erga, S. R. (2016). Chlorophyll a fluorescence lifetime reveals reversible UV-induced photosynthetic activity in the green algae *Tetraselmis*. European Biophysics Journal 45, 259–268. doi: 10.1007/s00249-015-1092-z

Ku, C., Sheyn, U., Sebé-Pedrós, A., Ben-Dor, S., Schatz, D., Tanay, A., et al. (2020). A single-cell view on alga-virus interactions reveals sequential transcriptional programs and infection states. Science Advances 6, eaba4137. doi: 10.1126/sciadv.aba4137

La Scola, B., Desnues, C., Pagnier, I., Robert, C., Barrassi, L., Fournous, G., et al. (2008). The virophage as a unique parasite of the giant mimivirus. Nature 455, 100–104. doi: 10.1038/nature07218

Lefkowitz, E. J., Dempsey, D. M., Hendrickson, R. C., Orton, R. J., Siddell, S. G., and Smith, D. B. (2018). Virus taxonomy: the database of the International Committee on Taxonomy of Viruses (ICTV). Nucleic Acids Research 46, D708–D717. doi: 10.1093/nar/gkx932

Liu, R., Liu, Y., Chen, Y., Zhan, Y., and Zeng, Q. (2019). Cyanobacterial viruses exhibit diurnal rhythms during infection. Proceedings of the National Academy of Sciences 116, 14077–14082. doi: 10.1073/pnas.1819689116

Mann, N. H., Cook, A., Millard, A., Bailey, S., and Clokie, M. (2003). Bacterial photosynthesis genes in a virus. Nature 424, 741–741. doi: 10.1038/424741a

Middelboe, M., and Brussaard, C. P. D. (2017). Marine viruses: Key players in marine ecosystems. Viruses 9, 302. doi: 10.3390/v9100302

Moniruzzaman, M., Gann, E. R., LeCleir, G. R., Kang, Y., Gobler, C. J., and Wilhelm, S. W. (2016). Diversity and dynamics of algal Megaviridae members during a harmful brown tide caused by the pelagophyte, *Aureococcus anophagefferens*. FEMS Microbiology Ecology 92, fiw058. doi: 10.1093/femsec/fiw058

Moniruzzaman, M., Gann, E. R., and Wilhelm, S. W. (2018). Infection by a Giant Virus (AaV) Induces widespread physiological reprogramming in *Aureococcus anophagefferens* CCMP1984 - A harmful bloom algae. Frontier in Microbiology 9, 752. doi: 10.3389/fmicb.2018.00752

Moniruzzaman, M., Martinez-Gutierrez, C. A., Weinheimer, A. R., and Aylward, F. O. (2020). Dynamic genome evolution and complex virocell metabolism of globally-distributed giant viruses. Nature Communications 11, 1710. doi: 10.1038/s41467-020-15507-2

Moniruzzaman, M., Wurch, L. L., Alexander, H., Dyhrman, S. T., Gobler, C. J., and Wilhelm, S. W. (2017). Virus-host relationships of marine single-celled eukaryotes resolved from metatranscriptomics. Nature Communications 8, 16054. doi: 10.1038/ncomms16054

Mulholland, M. R., Boneillo, G., and Minor, E. C. (2004). A comparison of N and C uptake during brown tide (*Aureococcus anophagefferens*) blooms from two coastal bays on the east coast of the USA. Harmful Algae 3, 361–376. doi: 10.1016/j.hal.2004.06.007

Ni, T., and Zeng, Q. (2016). Diel infection of cyanobacteria by cyanophages. Frontiers in Marine Science, 2, 123. doi: 10.3389/fmars.2015.00123

Nissimov, J. I., Talmy, D., Haramaty, L., Fredricks, H. F., Zelzion, E., Knowles, B., et al. (2019). Biochemical diversity of glycosphingolipid biosynthesis as a driver of *Coccolithovirus* competitive ecology. Environmental Microbiology 21, 2182–2197. doi: 10.1111/1462-2920.14633

Probyn, T. A., Bernard, S., Pitcher, G. C., and Pienaar, R. N. (2010). Ecophysiological studies on *Aureococcus anophagefferens* blooms in Saldanha Bay, South Africa. Harmful Algae 9, 123–133. doi: 10.1016/j.hal.2009.08.008

R Core Team (2021). R: A language and environment for statistical computing. Available at: https://www.R-project.org/.

Raoult, D., Audic, S., Robert, C., Abergel, C., Renesto, P., Ogata, H., et al. (2004). The 1.2-megabase genome sequence of Mimivirus. Science 306, 1344–1350. doi: 10.1126/science.1101485

Roux, S., Brum, J. R., Dutilh, B. E., Sunagawa, S., Duhaime, M. B., Loy, A., et al. (2016). Ecogenomics and potential biogeochemical impacts of globally abundant ocean viruses. Nature 537, 689–693. doi: 10.1038/nature19366

Rowe, J. M., Dunlap, J. R., Gobler, C. J., Anderson, O. R., Gastrich, M. D., and Wilhelm, S. W. (2008). Isolation of a non-phage-like lytic virus infecting *Aureococcus Anophagefferen*s. Journal of Phycology 44, 71–76. doi: 10.1111/j.1529-8817.2007.00453.x

Schulz, F., Roux, S., Paez-Espino, D., Jungbluth, S., Walsh, D. A., Denef, V. J., et al. (2020). Giant virus diversity and host interactions through global metagenomics. Nature 578, 432–436. doi: 10.1038/s41586-020-1957-x

Schvarcz, C. R., and Steward, G. F. (2018). A giant virus infecting green algae encodes key fermentation genes. Virology 518, 423–433. doi: 10.1016/j.virol.2018.03.010

Sheyn, U., Rosenwasser, S., Ben-Dor, S., Porat, Z., and Vardi, A. (2016). Modulation of host ROS metabolism is essential for viral infection of a bloom-forming coccolithophore in the ocean. ISME J 10, 1742–1754. doi: 10.1038/ismej.2015.228

Sieburth, J. McN., Johnson, P. W., and Hargraves, P. E. (1988). Ultrastructure and ecology of *Aureococcus anophageferens* Gen. Et Sp. Nov. (Chrysophyceae): The dominant picoplankter during a bloom in Narragansett Bay, Rhode Island, Summer 1985. Journal of Phycology 24, 416–425. doi: 10.1111/j.1529-8817.1988.tb04485.x

Sieracki, M. E., Gobler, C. J., Cucci, T. L., Thier, E. C., Gilg, I. C., and Keller, M. D. (2004). Pico- and nanoplankton dynamics during bloom initiation of *Aureococcus* in a Long Island, NY bay. Harmful Algae 3, 459–470. doi: 10.1016/j.hal.2004.06.012

Simjouw, J.-P., Mulholland, M. R., and Minor, E. C. (2004). Changes in dissolved organic matter characteristics in Chincoteague Bay during a bloom of the pelagophyte *Aureococcus anophagefferens*. Estuaries 27, 986–998. doi: 10.1007/BF02803425

Suttle, C. A., and Wilhelm, S. W. (1999). Viruses and nutrient cycles in the sea. BioScience 49. doi: 10.2307/1313569

Truchon, A. R., Chase, E. E., Gann, E. R., Moniruzzaman, M., Creasey, B. A., Aylward, F. O., et al. (2023). *Kratosvirus quantuckense*: The history and novelty of an algal bloom disrupting virus and a model for giant virus research. Frontiers in Microbiology 14.

Truchon, A. R., Chase, E. E., Stark, A. R., and Wilhelm, S. W. (2024). The diel disconnect between cell growth and division in *Aureococcus* is interrupted by giant virus infection. bioRxiv 2024.05.01.592014. doi: 10.1101/2024.05.01.592014

Truchon, A. R., Gann, E. R., and Wilhelm, S. W. (2022). Closed, circular genome sequence of Aureococcus anophagefferens Virus, a lytic virus of a brown tide-forming alga. Microbiology Resource Announcements 11, e00282–22. doi: 10.1128/mra.00282-22

Vincent, F., and Vardi, A. (2023). Viral infection in the ocean—A journey across scales. PLOS Biology 21, e3001966. doi: 10.1371/journal.pbio.3001966

Wazniak, C. E., and Glibert, P. M. (2004). Potential impacts of brown tide, *Aureococcus anophagefferens*, on juvenile hard clams, *Mercenaria mercenaria*, in the coastal bays of Maryland, USA. Harmful Algae 3, 321–329. doi: 10.1016/j.hal.2004.06.004

Wickham, H. (2011). ggplot2. WIREs Computational Statistics 3, 180–185. doi: 10.1002/wics.147

Wickham, H. (2023). stringr: Simple, Consistent Wrappers for Common String Operations. Available at: https://stringr.tidyverse.org

Wickham, H., Averick, M., Bryan, J., Chang, W., McGowan, L. D., François, R., et al. (2019). Welcome to the Tidyverse. Journal of Open Source Software 4, 1686. doi: 10.21105/joss.01686

Wickham, H., François, R., Henry, L., and Müller, K. (2022). dplyr: A Grammar of Data Manipulation. Available at: https://github.com/tidyverse/dplyr

Wickham, H., Vaughan, D., and Girlich, M. (2023). Tidyr: Easily tidy data with ‘spread()’ and ‘gather()’ functions. Available at: https://github.com/tidyverse/tidyr, https://tidyr.tidyverse.org

Xie, Z., Wu, Z., Wang, O., and Liu, F. (2023). Unexpected growth promotion of *Chlorella sacchrarophila* triggered by herbicides DCMU. Journal of Hazardous Materials 452, 131216. doi: 10.1016/j.jhazmat.2023.131216

Yao, P., Lei, L., Zhao, B., Wang, J., and Chen, L. (2019). Spatial-temporal variation of *Aureococcus anophagefferens* blooms in relation to environmental factors in the coastal waters of Qinhuangdao, China. Harmful Algae 86, 106–118. doi: 10.1016/j.hal.2019.05.011

Zhao, Y., Zhao, Y., Zheng, S., Zhao, L., Zhang, W., Xiao, T., et al. (2023). Enhanced resolution of marine viruses with violet side scatter. Cytometry Part A 103, 260–268. doi: 10.1002/cyto.a.24674

Zimmerman, A. E., Howard-Varona, C., Needham, D. M., John, S. G., Worden, A. Z., Sullivan, M. B., et al. (2020). Metabolic and biogeochemical consequences of viral infection in aquatic ecosystems. Nature Reviews Microbiology 18, 21–34. doi: 10.1038/s41579-019-0270-x

