## Supplemental Tables and Figures for "Time of day of infection shapes development of a eukaryotic algal-*Nucleocytoviricota* virocell"

For publication in conjunction with the following:

**FIGURES**

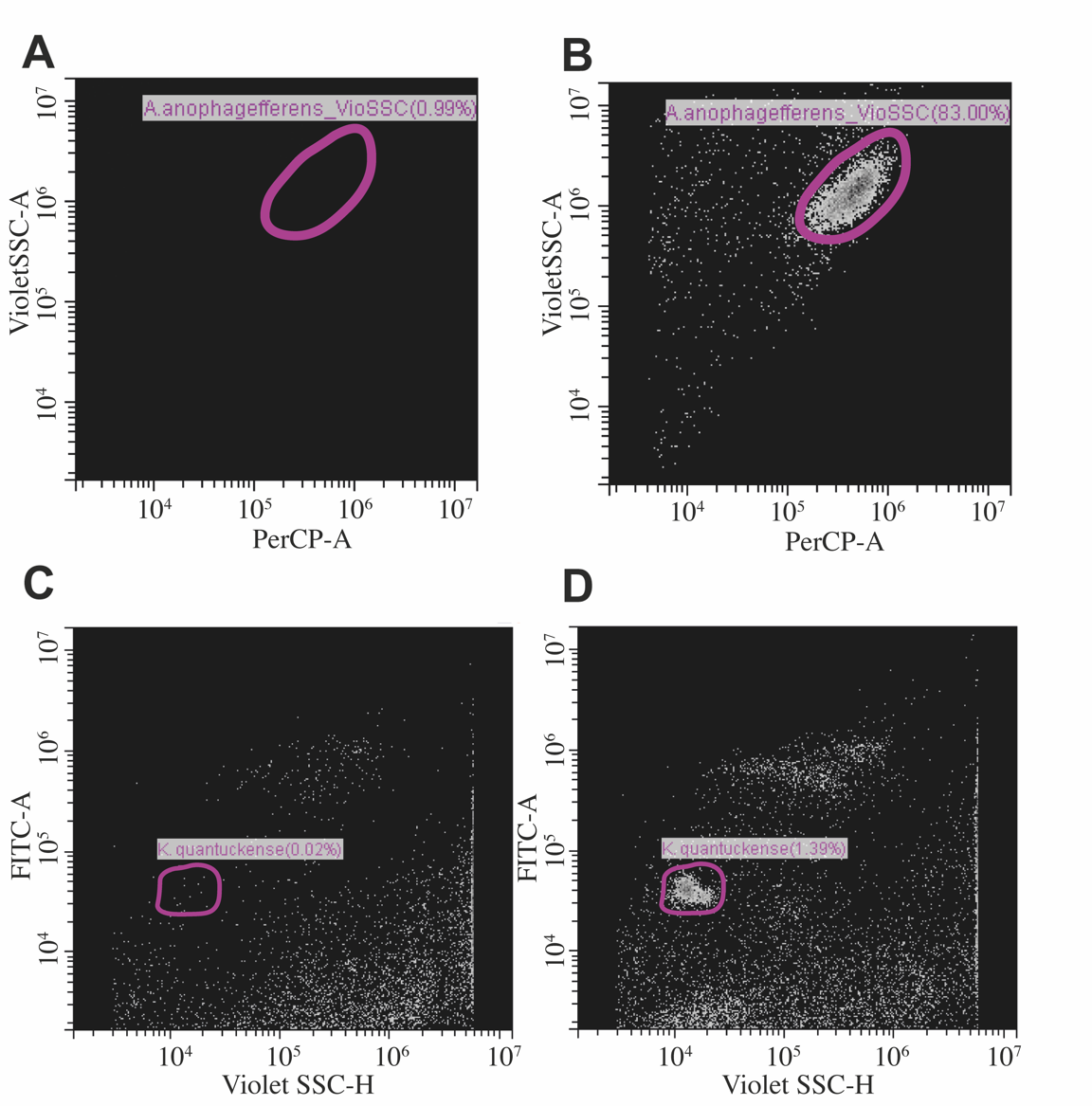

**Supplemental Figure 1.** Cytometry methodology for assessing *Aureococcus anophagefferens* cell populations and *Kratosvirus quantuckense* viral particles. (**A**) ASP12A run without *A. anophagefferens* culture as a negative control. (**B**) *A. anophagefferens* cells as located by PerCP background fluorescence and relative size (by Violet SSC). (**C**) Results of uninfected cultures stained with SYBR Gold and run using settings for detecting viruses. (**D**) *K. quantuckense* viral particles located using SYBR Gold staining and identified with FITC. (**A**) and (**B**) are run under the same gain settings (except (**A**) has a PerCP threshold of 1000 whereas (**B**) has a PerCP threshold of 5000) with the same population gated (or theoretical population area), additionally (**C**) and (**D**) have the same gain settings and the same population gated. PerCP; peridinin-chlorophyll protein complex, FITC-A; fluorescein isothiocyanate based on area of pulse, Violet SSC-H; side scatter based on height of violet laser pulse.

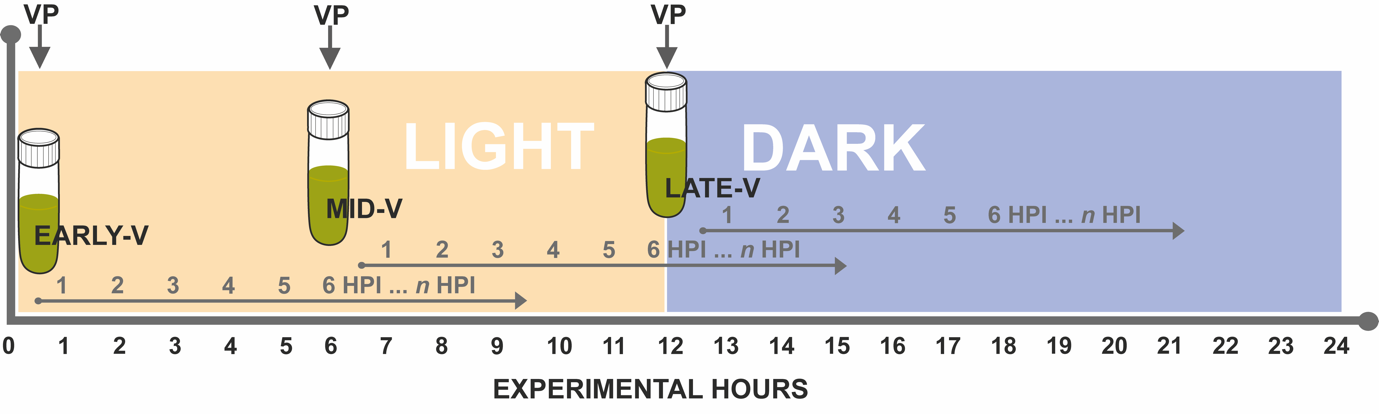

**Supplemental Figure 2.** Sampling schematic of virus lysate (*Kratosvirus quantuckense*) inoculation during the initial light cycle period (12 h). Experimental hours for the duration of full light/dark cycle (first 24 h) is shown, depicting when viral lysate was added to the EARLY-V (*i.e.,* 0 experimental hours), MID-V (*i.e.,* 6 experimental hours), and LATE-V (*i.e.,* ~20 minutes before 12 h) treatments. HPI; hours post infection, VP; viral particles (lysate), *n*; number of hours up until the end of the experiment (74 h).

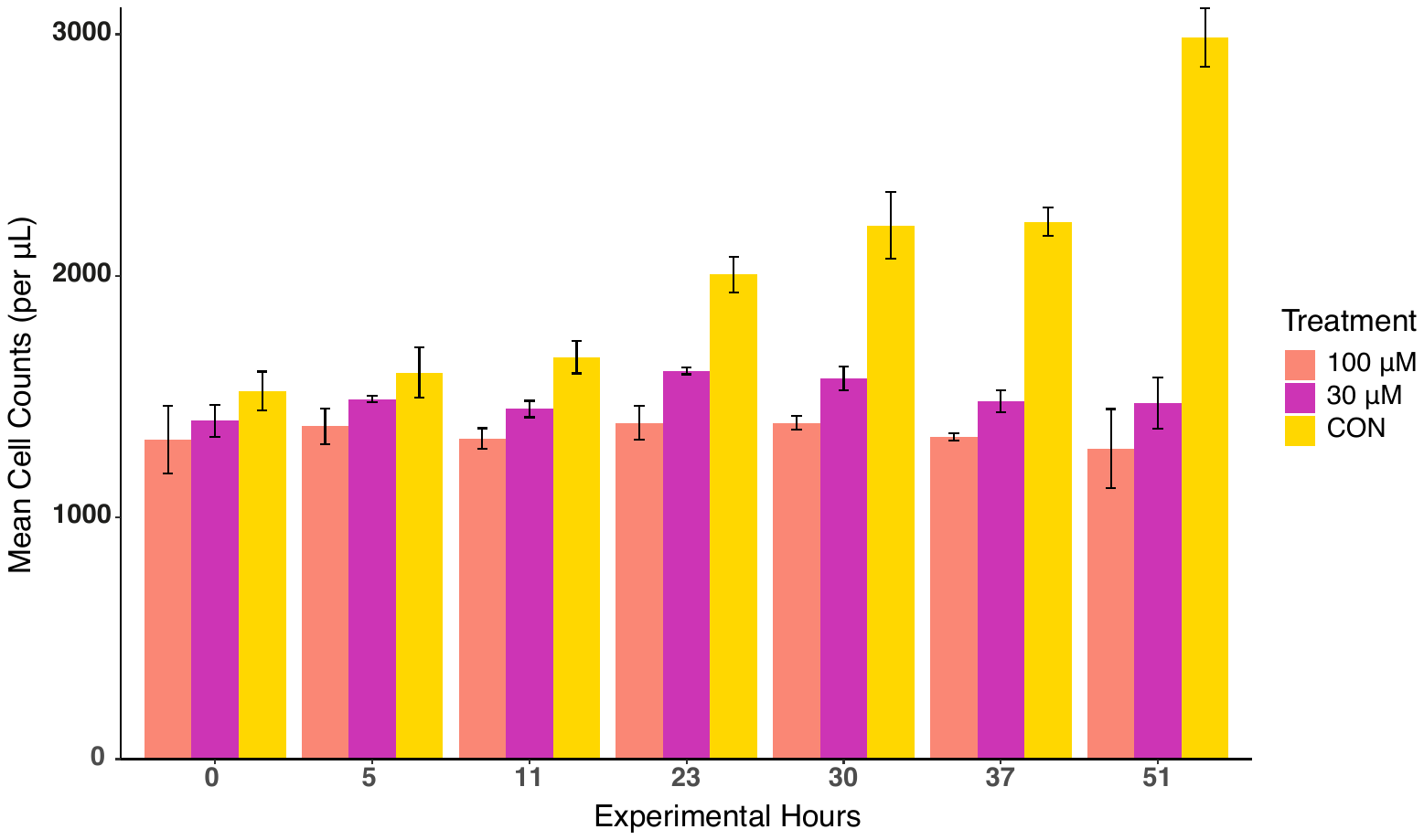

**Supplemental Figure 3.** *Aureococcus anophagefferens* cell counts over two days+ using flow cytometry to determine effective DCMU concentration for the DCMU treated *A. anophagefferens* infection (with *Kratosvirus quantuckense*). It was noted that the higher final concentration (100 μM) had fewer cells than the lower concentration (30 μM). CON; no DCMU applied.

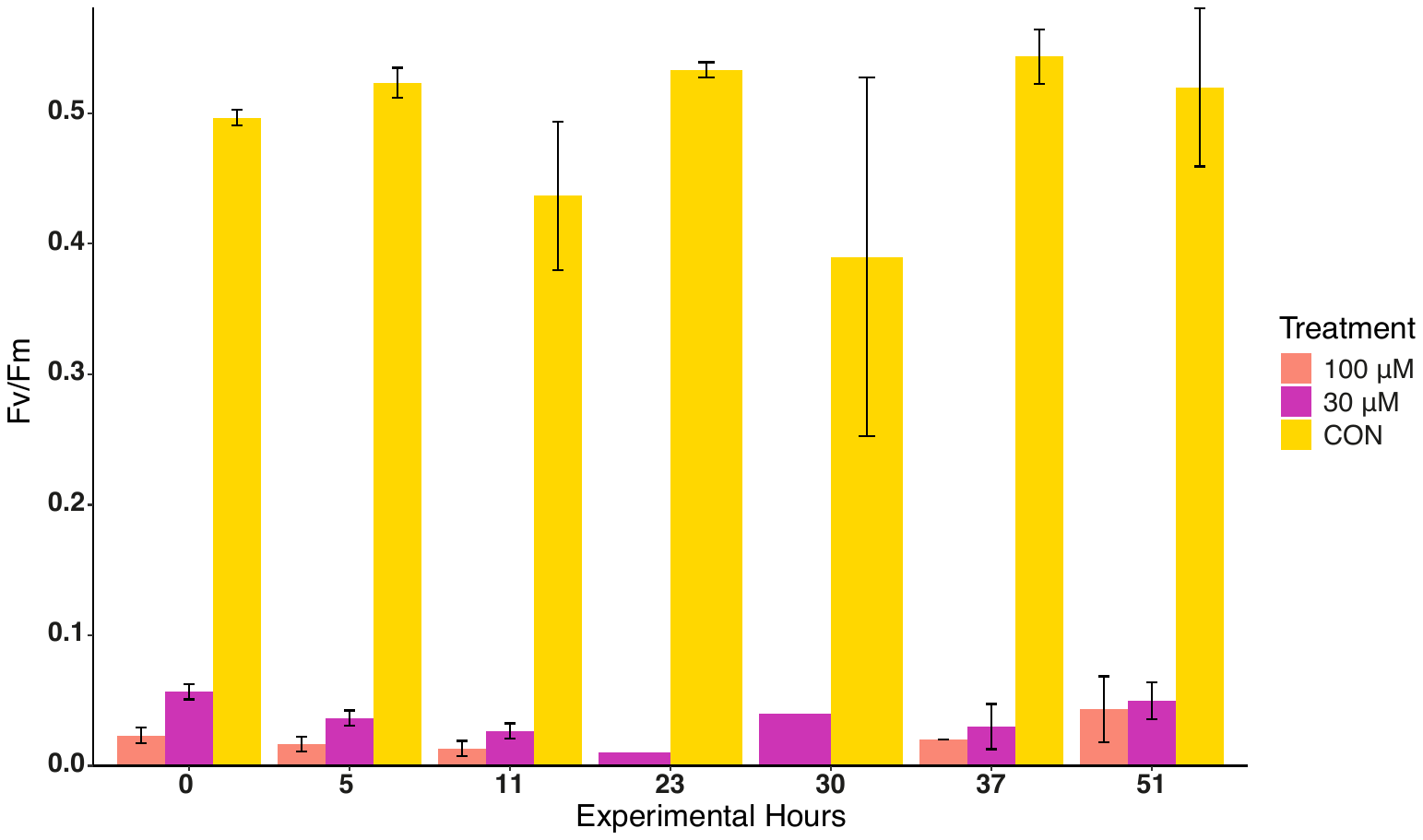

**Supplemental Figure 4.** *Aureococcus anophagefferens* F_v_/F_m_ over two days+ to determine DCMU concentration for a DCMU treated *A. anophagefferens* infection. It was noted that the higher final concentration (100 μM) of DCMU had a marginally lower F_v_/F_m_ (30 μM). CON; no DCMU applied. We determined a final concentration of 40 μM DCMU would reduce F_v_/F_m_ effectively while minimalizing the impacts on cell counts over the day cycle (24 h), given these results and **Supplemental Figure 3**.

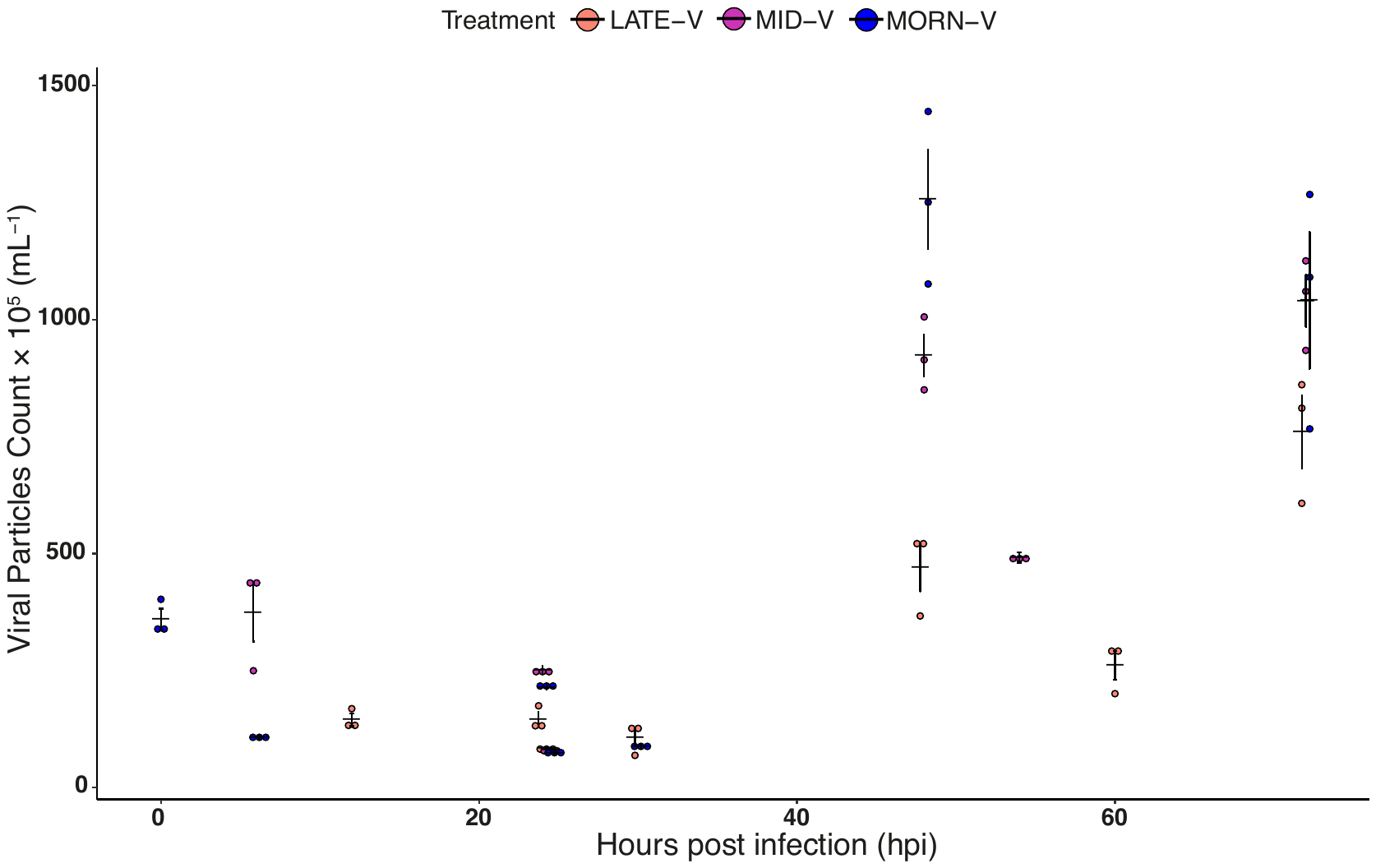

**Supplemental Figure 6.** Virocell formation (*i.e.,* virus lysate application) at different time points within a 12/12 light/dark cycle in *Aureococcus anophagefferens* with the virus *Kratosvirus quantuckense*. Mean viral particle counts (within culture media) at different time points (all hpi are relative to the early infection start time) on a linear time scale. LATE-V; lysate applied ~20 min before the end of the 12 h light cycle, MID-V; lysate applied midway through the light cycle (6 h), EARLY-V; lysate applied at the start of the light cycle (12 h exposure). This figure is in complement to **Figure 1A**.

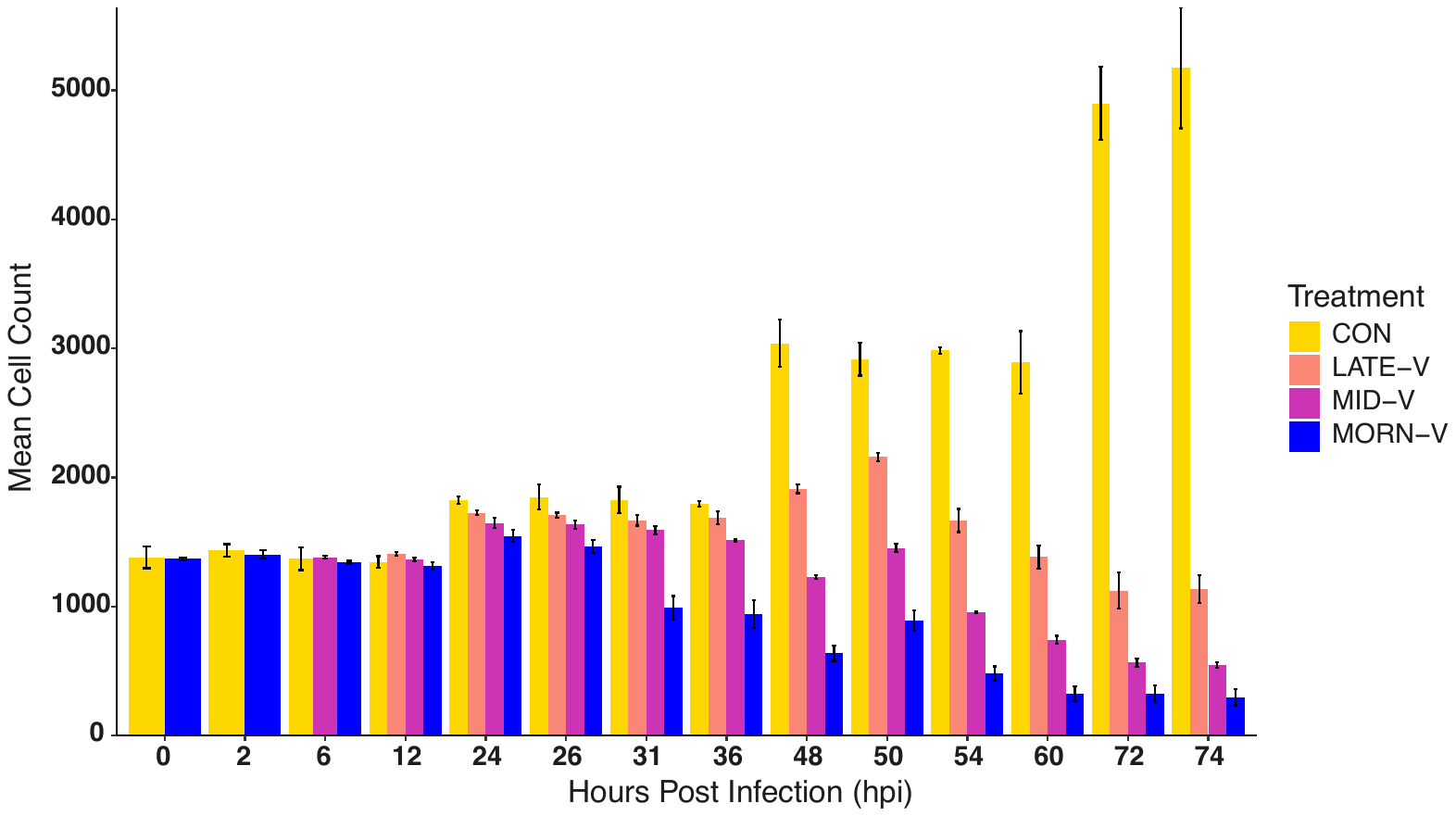

**Supplemental Figure 6.** Mean *Aureococcus anophagefferens* cell counts (per μL) using cytometry throughout the light exposure experiment. LATE-V; lysate applied ~20 min before the end of the 12 h light cycle, MID-V; lysate applied midway through the light cycle (6 h), EARLY-V; lysate applied at the start of the light cycle (12 h exposure), CON; ribocell control (no viral lysate).

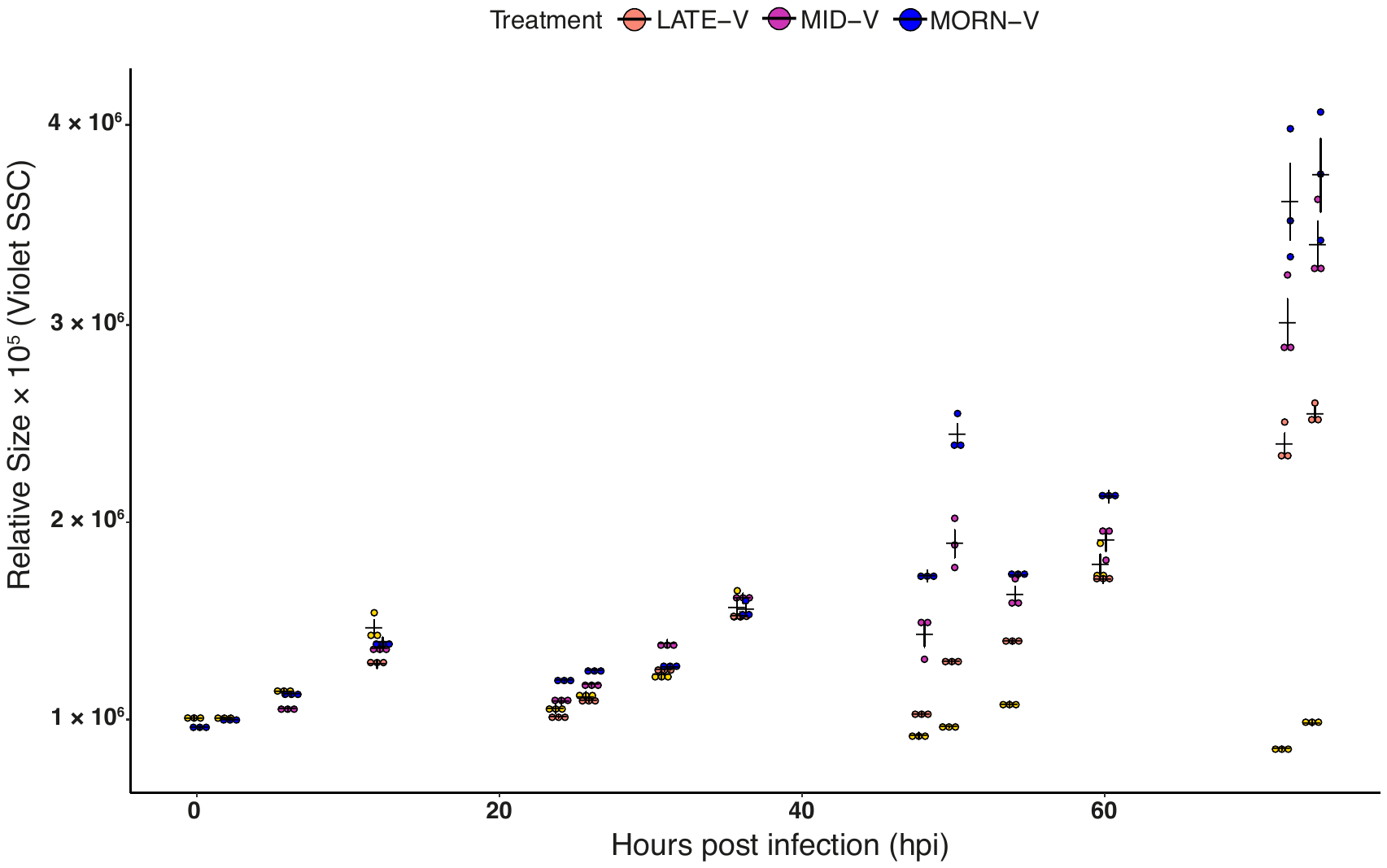

**Supplemental Figure 7.** Mean relative size of *Aureococcus anophagefferens* population cells throughout three light cycles (plus two hours after the third light cycle) on a linear time scale as measured via cytometry. LATE-V; lysate applied ~20 min before the end of the 12 h light cycle, MID-V; lysate applied midway through the light cycle (6 h), EARLY-V; lysate applied at the start of the light cycle (12 h exposure), CON; ribocell control (no viral lysate); SSC: side scatter. Violet SSC; side scatter using a violet laser. This figure is in complement to **Figure 1B**.

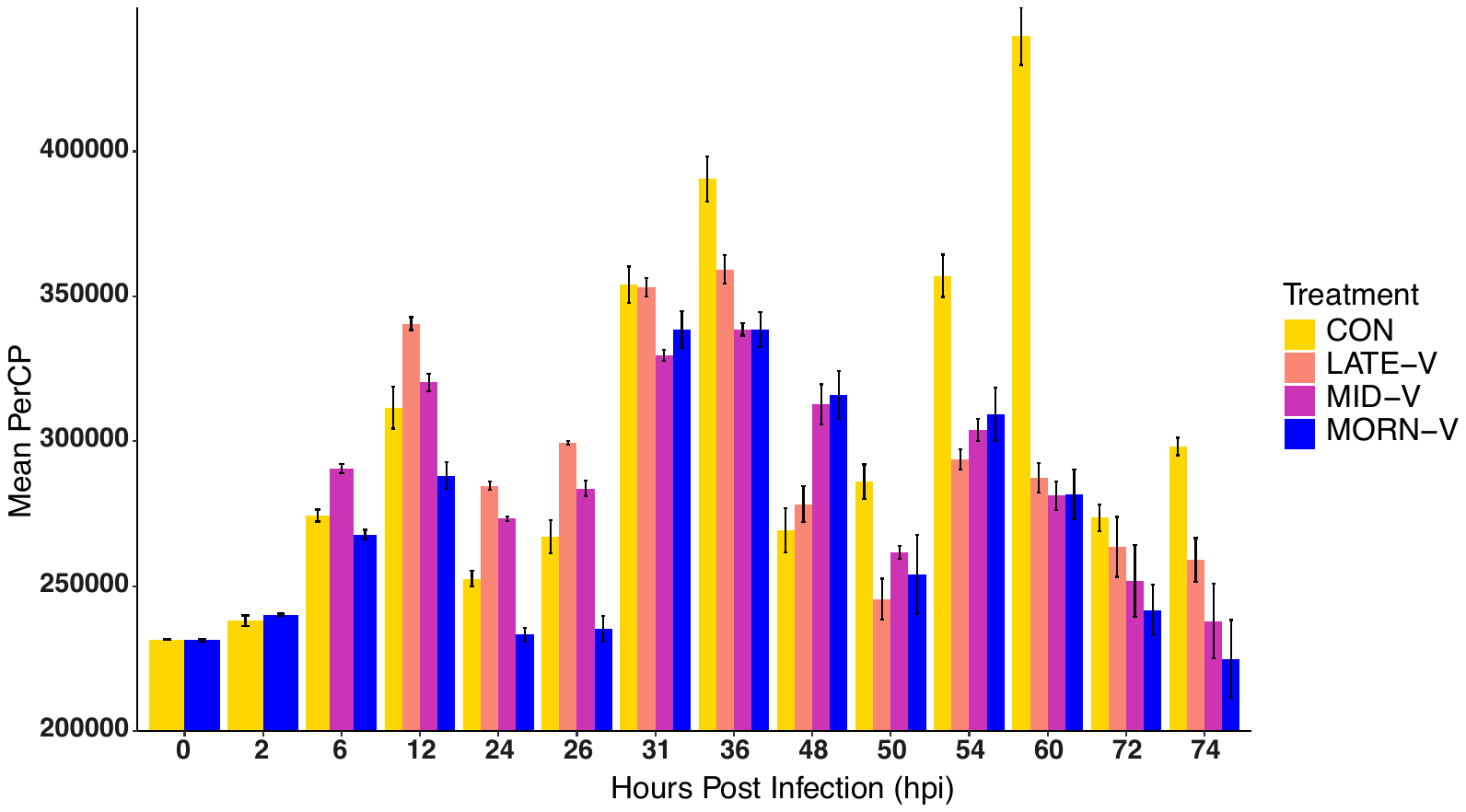

**Supplemental Figure 8.** Mean *Aureococcus anophagefferens* PerCP (Relative fluorescence units) using cytometry throughout the light exposure experiment. LATE-V; lysate applied ~20 min before the end of the 12 h light cycle, MID-V; lysate applied midway through the light cycle (6 h), EARLY-V; lysate applied at the start of the light cycle (12 h exposure), CON; ribocell control (no viral lysate).

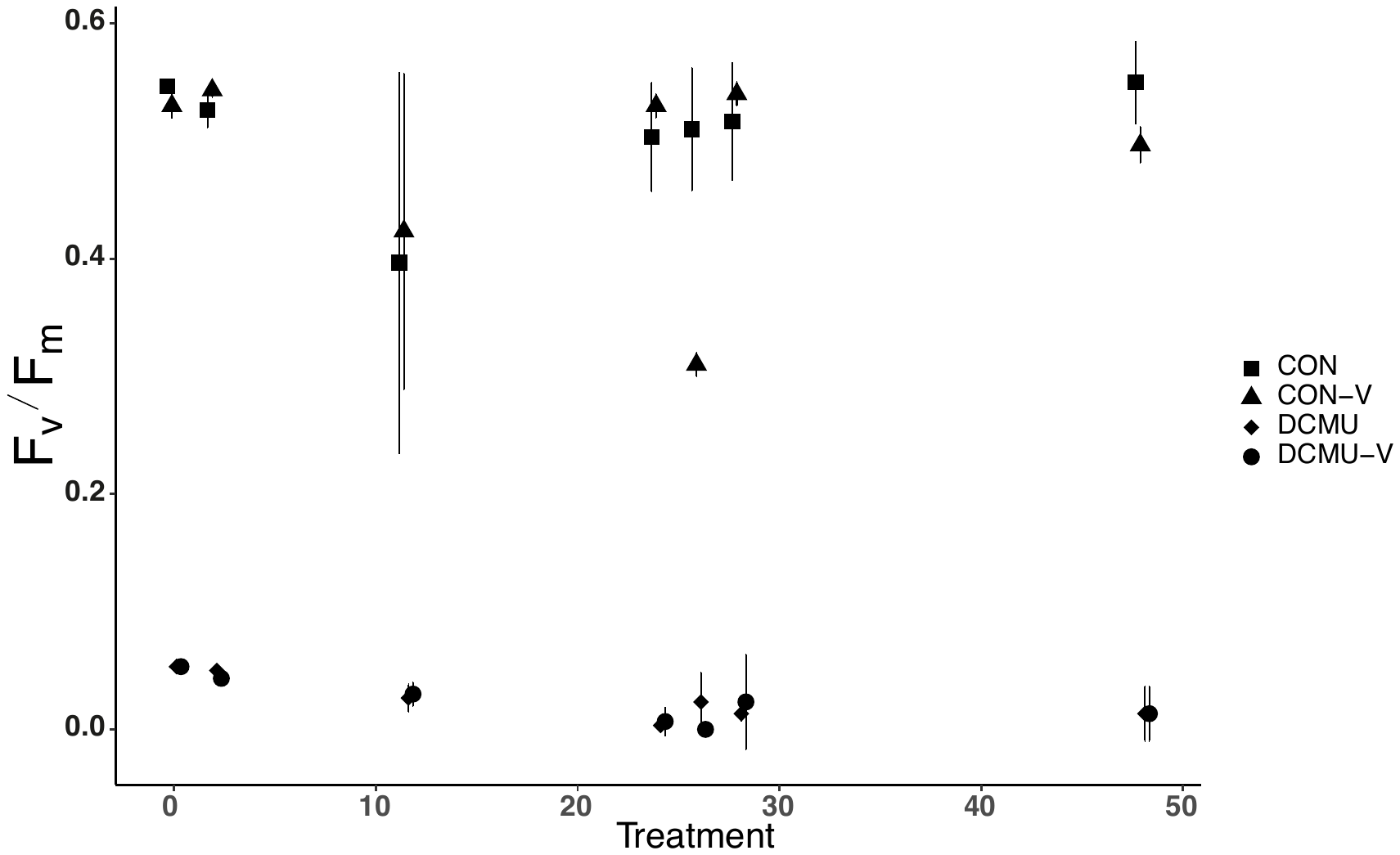

**Supplemental Figure 8.** *Aureococcus anophagefferens* photochemical response (F_v_/F_m_; variable to maximal fluorescence) to application of 40 μM DCMU (3-(3,4-dichlorophenyl)-1,1-dimethylurea; diuron) and infection by *Kratosvirus quantuckense* over two light cycles on a linear time scale. CON-V; virocells without DCMU applied, DCMU; ribocells with DCMU applied, DCMU-V; virocells with DCMU applied. This figure is in complement to **Figure 2A**.

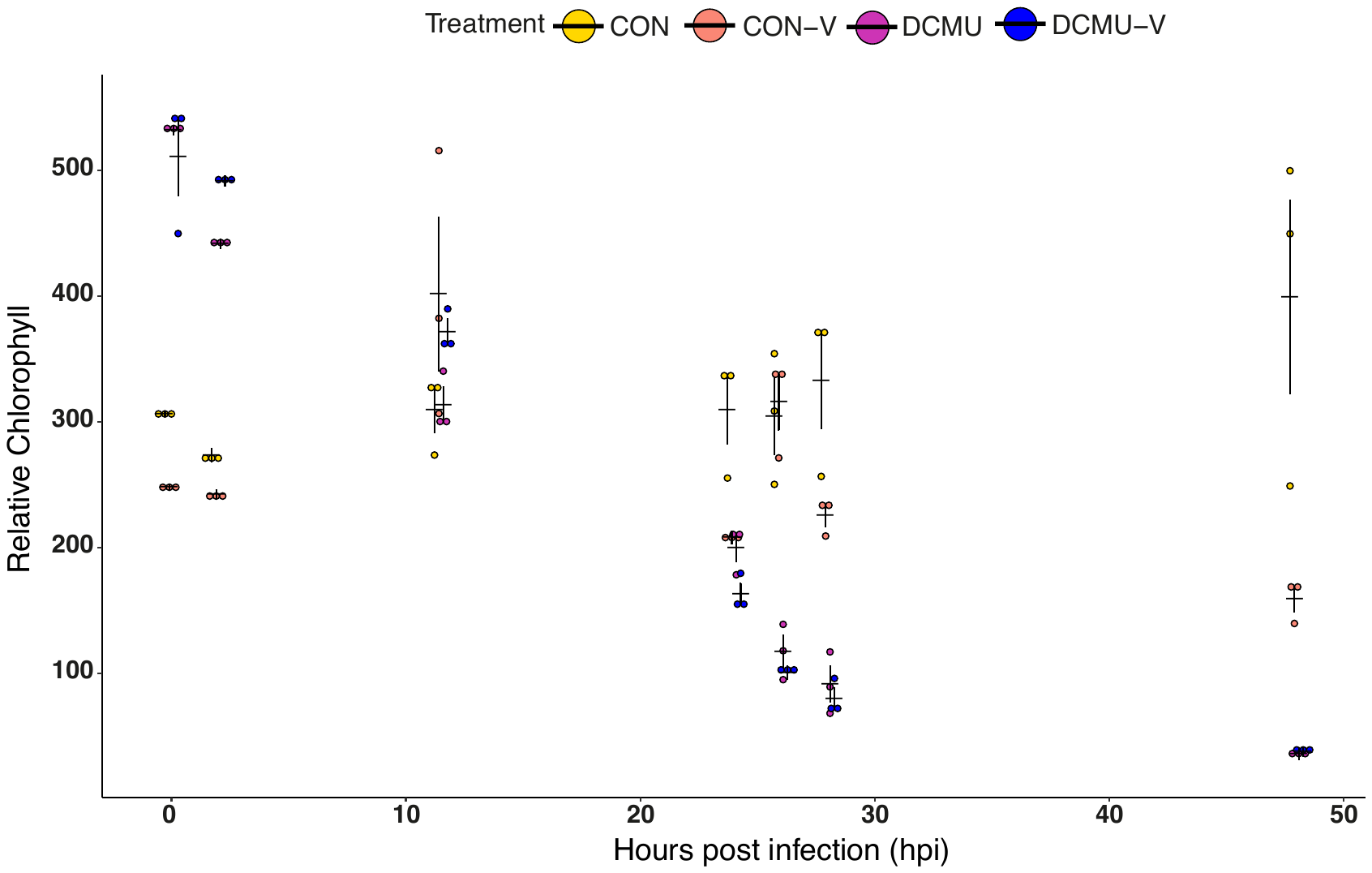

**Supplemental Figure 9.** Mean relative chlorophyll (by photochemistry) of *Aureococcus anophagefferens* population cells in the DCMU infection experiment on a linear time scale. CON; cells without DCMU or viral lysate applied, CON-V; virocells without DCMU applied, DCMU; ribocells with DCMU applied, DCMU-V; virocells with DCMU applied. This figure is in complement to **Figure 2B**.

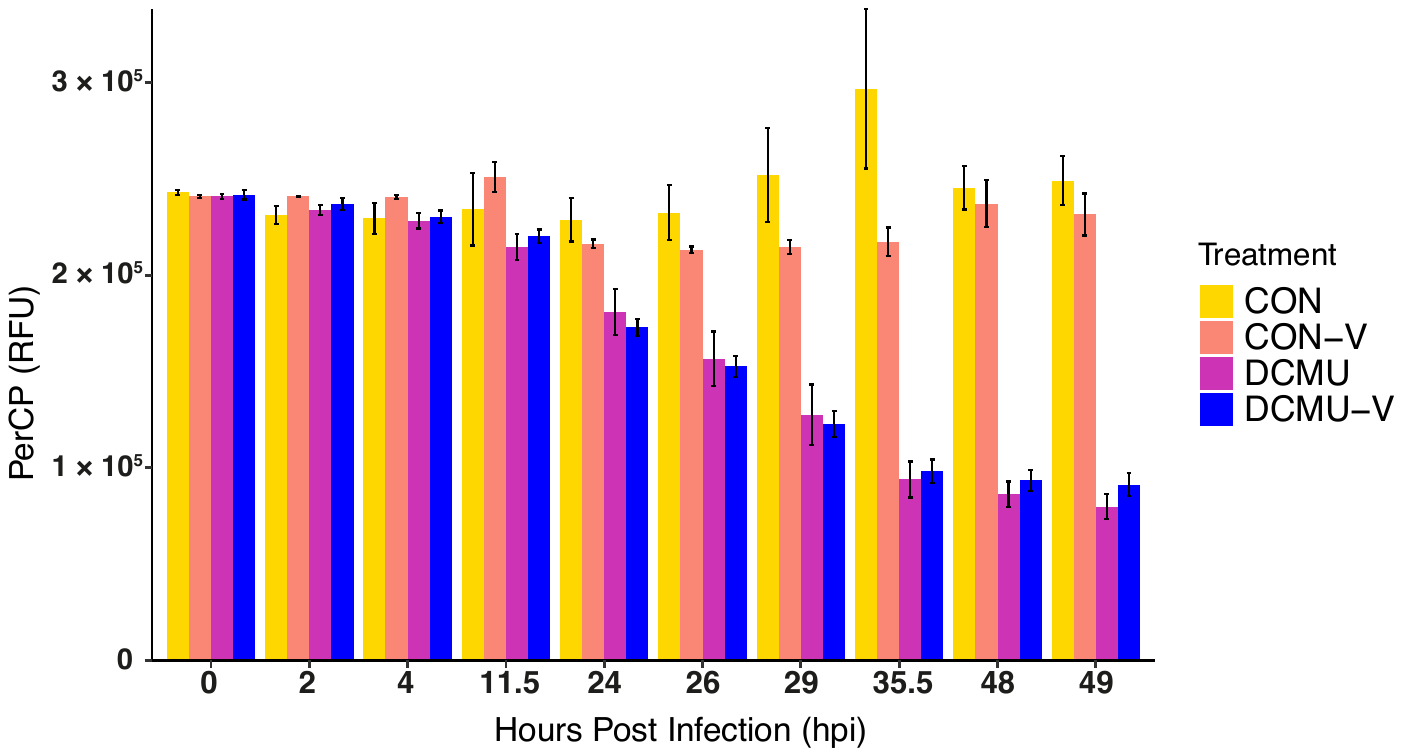

**Supplemental Figure 10.** Mean PerCP (by cytometry) of *Aureococcus anophagefferens* cells within the DCMU infection experiment. PerCP; peridinin-chlorophyll protein complex, RFU; relative fluorescence units, CON; no virus or DCMU applied, CON-V; virocells without DCMU applied, DCMU; ribocells with DCMU applied, DCMU-V; virocells with DCMU applied.

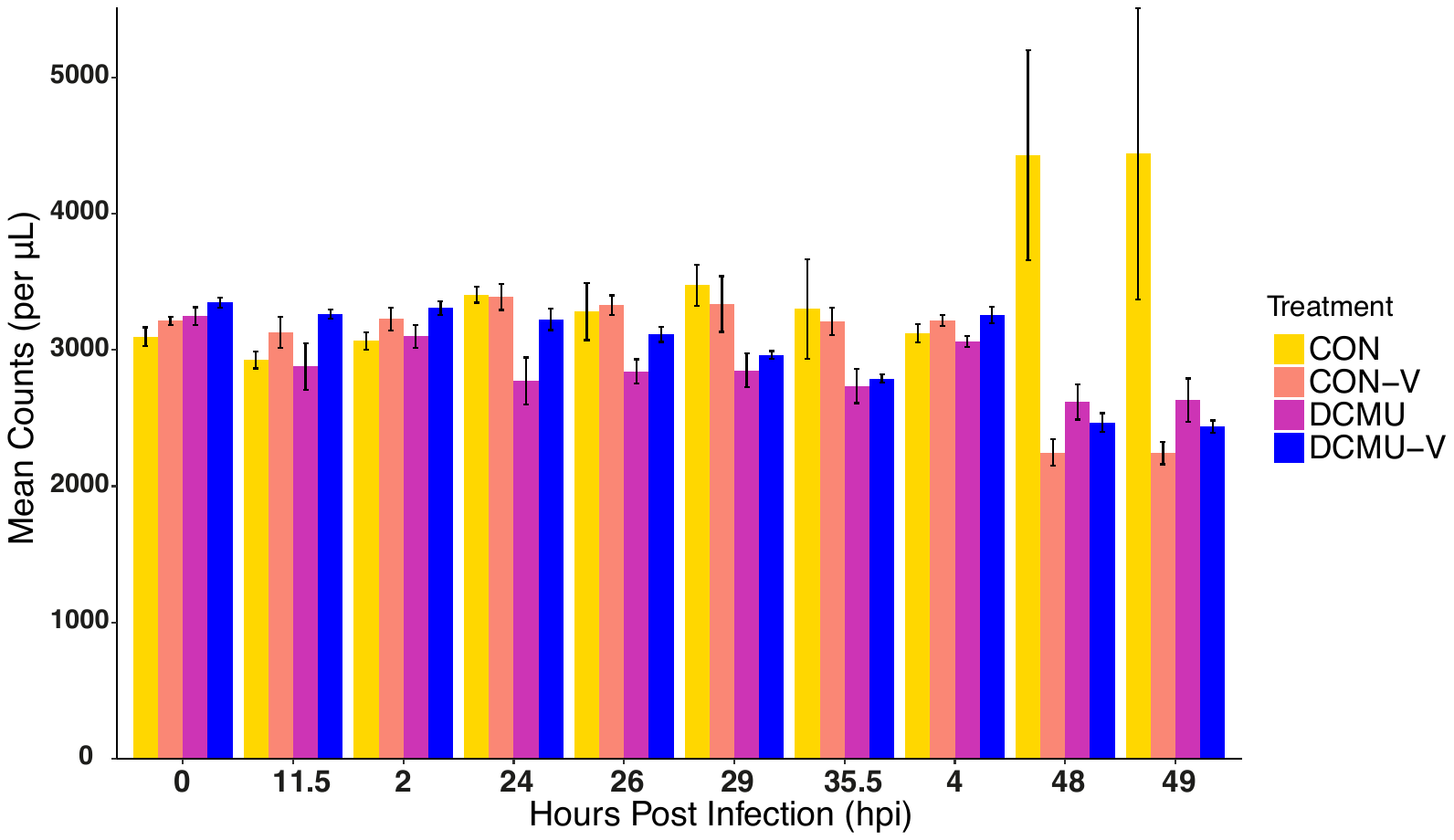

**Supplemental Figure 11.** Mean *Aureococcus anophagefferens* cell counts (per μL) using cytometry throughout the DCMU infection experiment. CON; no virus or DCMU applied, CON-V; virocells without DCMU applied, DCMU; ribocells with DCMU applied, DCMU-V; virocells with DCMU applied.

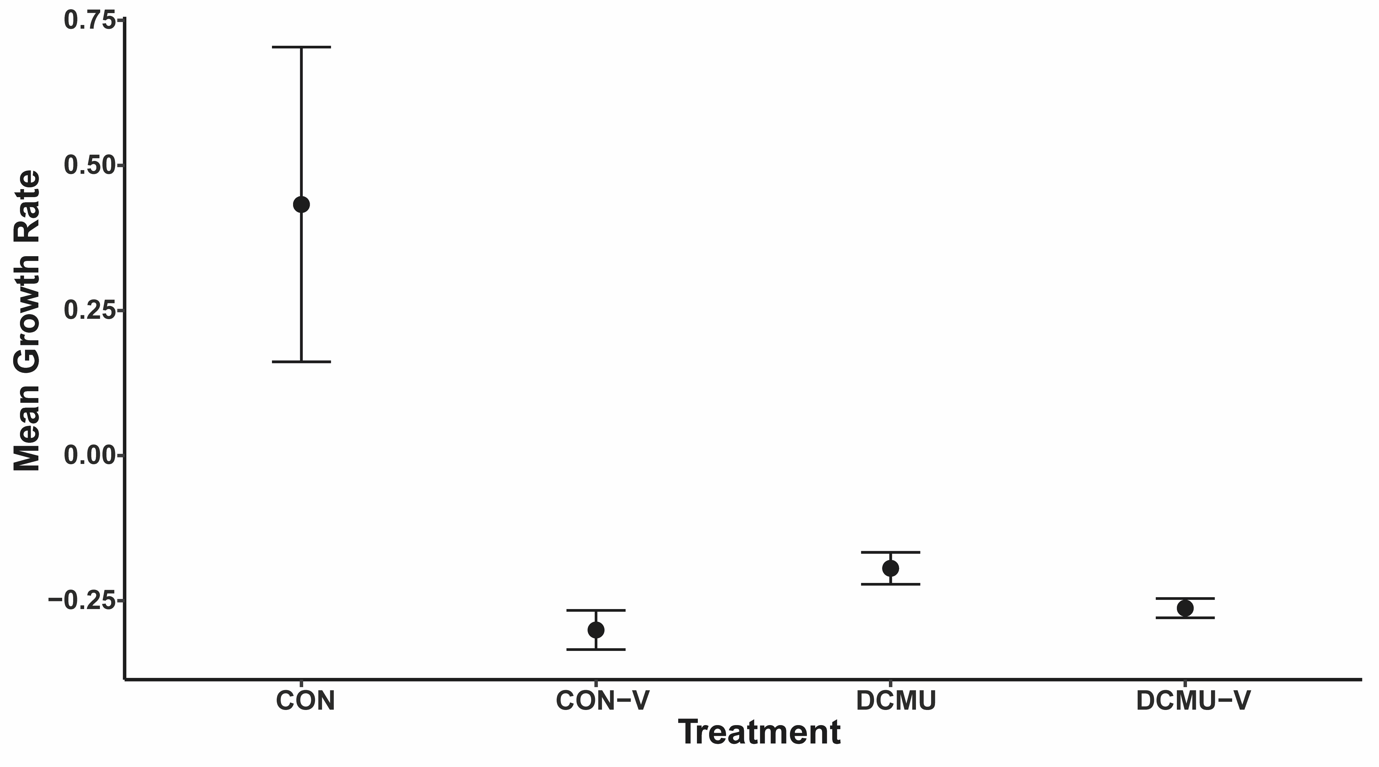

**Supplemental Figure 12.** *Aureococcus anophagefferens* growth rates (via cell population counts) per treatment within the DCMU experiment from 0 to 48 hpi (12/12 light and dark cycle). CON; control (ribocells), CON-V; virocells without DCMU applied, DCMU; ribocells with DCMU applied, DCMU-V; virocells with DCMU applied.

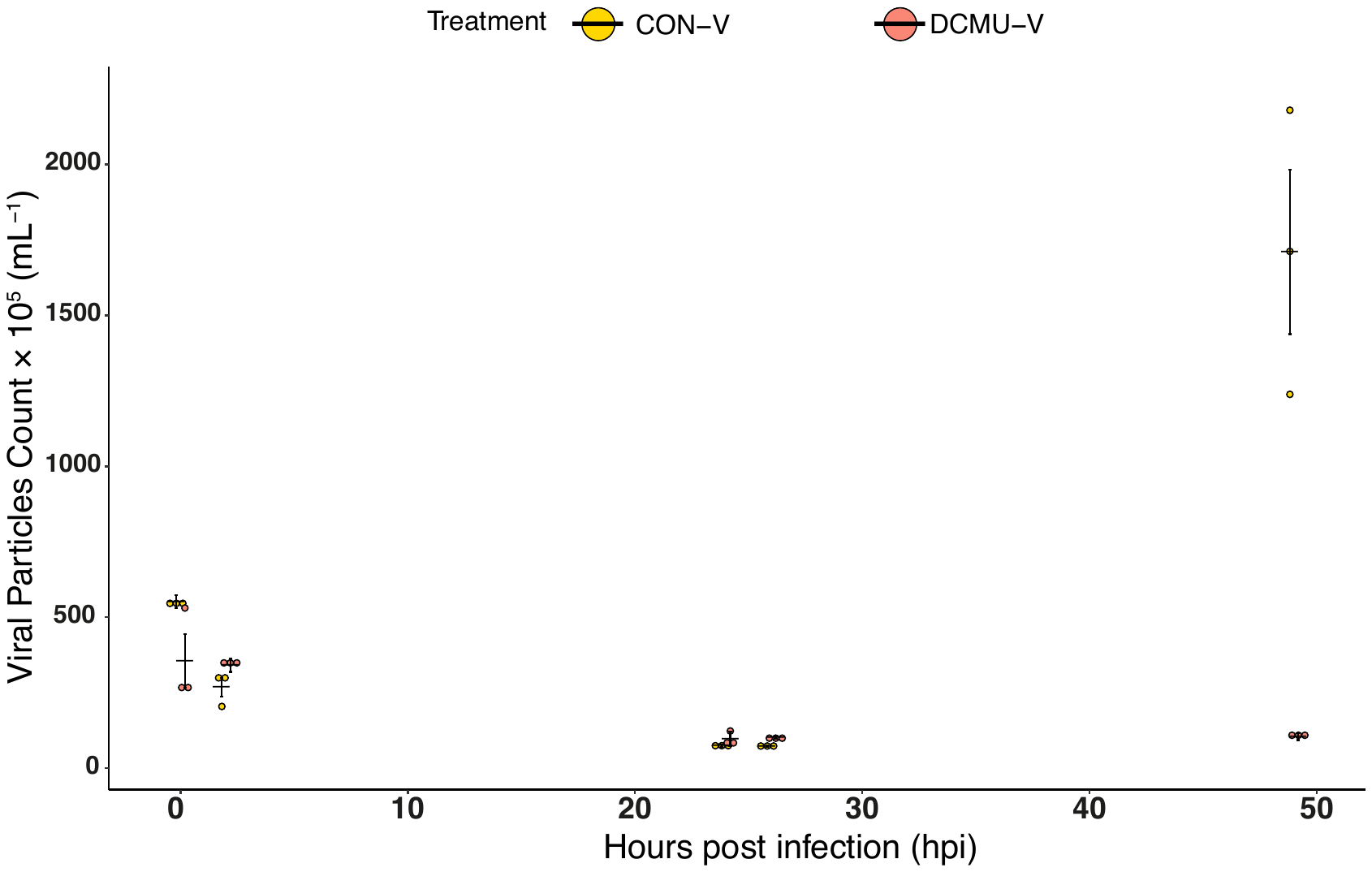

**Supplemental Figure 3.** Mean viral particle counts (within culture media) at different time points for the DCMU infection experiment on a linear time scale. This figure is in complement to **Figure 2C**.

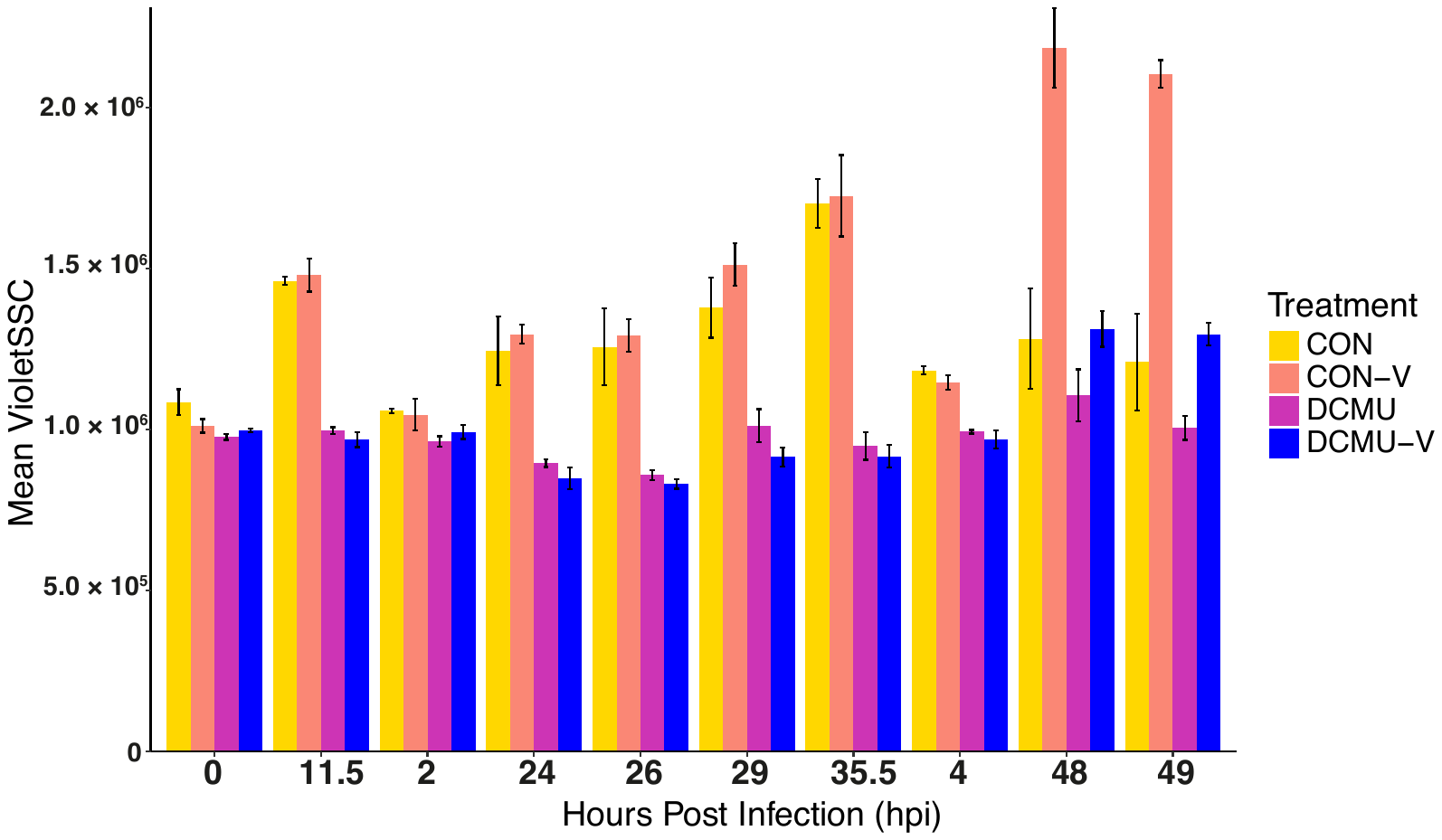

**Supplemental Figure 14.** Mean relative size of *Aureococcus anophagefferens* population cells throughout throughout 2+ light/dark cycles on a linear time scale as measured via cytometry. SSC: side scatter. Violet SSC; side scatter using a violet laser, CON-V; virocells without DCMU applied, DCMU; ribocells with DCMU applied, DCMU-V; virocells with DCMU applied.

**TABLES**

| **Supplemental Table 1.** Welch two sample *t*-test of viral counts across treatments in the varying light exposure infection experiment. A bold p-value indicates comparisons that were considered significant. Trt1; treatment 1, Trt2; treatment 2, *t*; test statistic, df; degrees of freedom. α = 0.05 | | | | | |
| --- | --- | --- | --- | --- | --- |
| **hpi** | **Trt1** | **Trt2** | ***t*** | **df** | **p-value** |
| 48 | LATE -V | MID-V | -6.6067 | 3.9307 | **0.0029** |
| 48 | EARLY-V | MID-V | 2.8874 | 2.6986 | **0.0717** |
| 48 | EARLY-V | LATE-V | 2.8926 | 2.8926 | **0.0077** |
| 72 | LATE -V | MID-V | -2.9291 | 3.647 | **0.0480** |
| 72 | EARLY-V | MID-V | -0.0091044 | 2.5743 | 0.9934 |
| 72 | EARLY-V | LATE-V | 1.8067 | 8.0623 | 0.1082 |

| **Supplemental Table 2.** Welch two sample *t*-test of violet side scatter (relative size) across treatments in the varying light exposure infection experiment. A bold p-value indicates comparisons that were considered significant. Trt1; treatment 1, Trt2; treatment 2, *t*; test statistic, df; degrees of freedom. α = 0.05 | | | | | |
| --- | --- | --- | --- | --- | --- |
| **hpi** | **Trt1** | **Trt2** | ***t*** | **df** | **p-value** |
| 24 | EARLY-V | LATE-V | -6.0873 | 2.3595 | **0.0172** |
| 24 | MID-V | LATE-V | -4.4414 | 2.8214 | **0.0241** |
| 24 | EARLY-V | MID-V | 2.6652 | 3.4323 | 0.0659 |
| 24 | CON | EARLY-V | 5.9139 | 2.4677 | **0.0164** |
| 24 | CON | MID-V | 1.3264 | 3.2766 | 0.2696 |
| 24 | CON | LATE-V | 1.7956 | 2.3451 | 0.1955 |
| 48 | EARLY-V | LATE-V | -23.985 | 2.2407 | **0.0010** |
| 48 | MID-V | LATE-V | -6.3899 | 2.0495 | **0.0222** |
| 48 | EARLY-V | MID-V | 4.3335 | 2.7874 | **0.0263** |
| 48 | CON | EARLY-V | 23.539 | 3.5414 | **5.098e-05** |
| 48 | CON | MID-V | 7.804 | 2.3827 | **0.0095** |
| 48 | CON | LATE-V | 5.2799 | 2.5048 | **0.0206** |
| 72 | EARLY-V | LATE-V | -17.927 | 3.9073 | **6.736e-05** |
| 72 | MID-V | LATE-V | -5.1109 | 2.9111 | **0.0156** |
| 72 | EARLY-V | MID-V | 2.6652 | 3.4323 | 0.0659 |
| 72 | CON | EARLY-V | 14.308 | 2.0147 | **0.0047** |
| 72 | CON | MID-V | 17.104 | 2.0347 | **0.0032** |
| 72 | CON | LATE-V | 25.979 | 2.1615 | **0.0010** |

| **Supplemental Table 3.** Welch two sample *t*-test of F_v_/F_m_ (at 40 min) of cells (*Aureococcus anophagefferens*) with and without DCMU application. A bold p-value indicates comparisons that were considered significant. Trt1; treatment 1, Trt2; treatment 2, *t*; test statistic, df; degrees of freedom. α = 0.05 | | | | |
| --- | --- | --- | --- | --- |
| **Trt1** | **Trt2** | ***t*** | **df** | **p-value** |
| CON | DCMU | -104.65 | 4 | **4.999e-08** |

| **Supplemental Table 4.** Welch two sample *t*-test on control and DCMU applied cells’ (*Aureococcus anophagefferens*) relative chlorophyll content (photochemistry). A bold p-value indicates comparisons that were considered significant. Trt1; treatment 1, Trt2; treatment 2, *t*; test statistic, df; degrees of freedom. α = 0.05 | | | | | |
| --- | --- | --- | --- | --- | --- |
| **hpi** | **Trt1** | **Trt2** | ***t*** | **df** | **p-value** |
| 0 | CON | DCMU | 56.953 | 2.967 | **1.322e-05** |
| 2 | CON | DCMU | 28.595 | 3.5087 | **2.772e-05** |
| 11.5 | CON | DCMU | 0.18862 | 3.7574 | 0.8601 |

| **Supplemental Table 5.** Welch two sample *t*-test of cell (*Aureococcus anophagefferens*) counts with the DCMU infection experiment. A bold p-value indicates comparisons that were considered significant. Trt1; treatment 1, Trt2; treatment 2, *t*; test statistic, df; degrees of freedom.. α = 0.05 | | | | | |
| --- | --- | --- | --- | --- | --- |
| **hpi** | **Trt1** | **Trt2** | ***t*** | **df** | **p-value** |
| 24 | CON | DCMU | -6.0305 | 2.4672 | **0.0156** |
| 26 | CON | DCMU | -3.3475 | 2.6942 | 0.0517 |
| 29 | CON | DCMU | -5.549 | 3.8516 | **0.0058** |
| 48 | CON | DCMU | -4.0051 | 2.1116 | 0.0521 |
| 48 | CON | CON-V | -4.8572 | 2.0608 | **0.0376** |
| 48 | CON | DCMU-V | -4.383 | 2.0325 | **0.0470** |
| 48 | CON-V | DCMU | 4.0131 | 3.6791 | **0.0188** |
| 48 | DCMU | DCMU-V | 1.7911 | 3.072 | 0.1690 |

| **Supplemental Table 6.** Welch two sample *t*-test of cell (*Aureococcus anophagefferens*) growth rates over 48 h with the DCMU infection experiment. A bold p-value indicates comparisons that were considered significant. Trt1; treatment 1, Trt2; treatment 2, *t*; test statistic, df; degrees of freedom. α = 0.05 | | | | |
| --- | --- | --- | --- | --- |
| **Trt1** | **Trt2** | ***t*** | **df** | **p-value** |
| CON | DCMU | 3.9835 | 2.0412 | 0.0557 |
| CON | DCMU-V | 4.4343 | 2.015 | **0.0466** |
| CON | CON-V | -4.6472 | 2.0617 | **0.0409** |
| CON-V | DCMU | -4.2339 | 3.8478 | **0.0145** |
| CON-V | DCMU-V | -1.7357 | 2.9184 | 0.1836 |
| DCMU | DCMU-V | -3.6996 | 3.2854 | **0.0294** |

| **Supplemental Table 7.** Welch two sample *t*-test of viral counts (at 48 hpi) of infected cells with and without DCMU applied. A bold p-value indicates comparisons that were considered significant. Trt1; treatment 1, Trt2; treatment 2, *t*; test statistic, df; degrees of freedom. α = 0.05 (values slightly higher were considered nearly significant). | | | | |
| --- | --- | --- | --- | --- |
| **Trt1** | **Trt2** | ***t*** | **df** | **p-value** |
| CON-V | DCMU-V | 5.9044 | 2.0068 | **0.02729** |
